# Rapid Production and Cryo-EM Structure Determination of GPCR-Nanodisc Complexes via Cell-Free Synthesis

**DOI:** 10.64898/2026.10.21.760646

**Authors:** Thomas A. Guilliam, Agata K. Jakimowicz, Marco M. Manni, Zachary B. Gaber, Giulia Paris, Steven W. Hardwick, Lee Cooper, Dimitri Y. Chirgadze, Tobias W.B. Ost

## Abstract

G protein-coupled receptors (GPCRs) represent major therapeutic targets, yet cryo-EM structure determination of unliganded or antagonist-bound GPCRs remains severely bottlenecked by small receptor mass, detergent-induced instability, and multi-week cell-based production pipelines. Here, we combine intracellular loop 3 (ICL3) rigid fiducial engineering with a high-throughput, cell-free protein synthesis workflow to rapidly generate cryo-EM-ready GPCR–nanodisc assemblies. Using the eProtein Discovery™ digital microfluidics system, we performed automated nanoliter-scale screening of construct fusions and nanodisc lipid compositions, establishing that co-translational insertion into nanodiscs is strictly required for receptor solubilization and affinity purification. Direct scale-up of screening conditions yielded structural-grade protein within four days from gene assembly to grid vitrification. We demonstrate this workflow with a 3.38 Å cryo-EM structure of antagonist-bound human adenosine A_2A_ receptor (A_2A_R)-β-lactamase fusion in complex with β-lactamase inhibitory protein II (BLIP-II). To demonstrate generalizability of protein quality, we extended this detergent-free co-translational pipeline to an expanded panel of 12 wild-type GPCRs spanning Classes A, B, and C, as well as six orphan receptors. All targets were purified with biophysical quality comparable to the A_2A_R benchmark. By removing the reliance on cellular expression or agonist/G-protein stabilization and compressing construct assembly, nanodisc screening, biophysical quality control, and grid vitrification into a four-day turnaround, this detergent-free cell-free pipeline provides a rapid Design–Build–Test–Learn (DBTL) engine to accelerate structure-based drug discovery for challenging and unliganded GPCR targets.

## Introduction

G protein-coupled receptors (GPCRs) represent the largest class of membrane proteins in the human genome and are targeted by approximately one-third of all FDA-approved therapeutics (Santos *et al*., 2017). Consequently, high-resolution structure determination of GPCRs is essential for deciphering receptor mechanisms and driving structure-based drug design.

In recent years, single-particle cryogenic-electron microscopy (cryo-EM) has revolutionized protein structure determination and removed the hurdle of protein crystallization which was a prerequisite for traditional X-ray crystallography approaches (Cheng, 2015). However, significant challenges remain when applying cryo-EM to GPCRs. GPCRs are relatively small proteins (∼40-50 kDa) and their distinguishing structural architecture is embedded within the cell membrane. These characteristics make particle alignment for cryo-EM extremely difficult. To circumvent these issues, most cryo-EM approaches focus on GPCR-signaling protein complexes, further stabilized with antibodies to provide rigid structural features to aid particle alignment (Zhang *et al*., 2022). Unfortunately, this approach does not enable structure determination in the absence of signaling proteins as is required for antagonist-bound or unliganded GPCR conformations (Zhang *et al*., 2022). Similarly, obtaining signaling proteins and forming a stable complex with the GPCR can prove challenging, further slowing the path to a high-resolution structure. The few cryo-EM structures of unliganded GPCRs that do exist are typically of receptors that form stable dimers or have rigid extracellular domains (Du *et al*., 2021; Josephs *et al*., 2021; Koehl *et al*., 2019).

Since GPCRs share a common domain architecture (Rosenbaum *et al*., 2009), an alternative broadly applicable approach is to engineer a rigid fusion protein between transmembrane (TM) helices 5 and 6 to replace the third intracellular loop (ICL3). This strategy was classically used during X-ray crystallography to form crystal contacts via the fusion partner (Chun *et al*., 2012; Rosenbaum *et al*., 2007). Recently, fusion proteins in ICL3 have been utilized as rigid fiducials to aid particle alignment during cryo-EM (Singh *et al*., 2026). Generally, an additional high-affinity antibody or binding partner to the fusion protein is included to add mass and improve resolution. This strategy is exemplified in a recent paper by Shah *et al*. where a 3.2 Å resolution structure of insect cell-expressed and detergent-solubilized antagonist-bound adenosine receptor A_2A_ (A_2A_R) was obtained by fusing β-lactamase in ICL3 and forming a complex with β-lactamase inhibitory protein II (BLIP-II) (Shah *et al*., 2026). A similar approach has recently been used to achieve a 4.2 Å resolution structure of ultra-thermostable β1-adrenoceptor containing a β-lactamase fusion in ICL3 (Collu *et al*., 2026).

While this strategy solves some of the issues associated with high-resolution cryo-EM of GPCRs, obtaining sufficient mass of high-quality protein from cellular expression systems is slow and can require several rounds of multi-week optimization. Moreover, purification of GPCRs expressed in cells requires harsh detergent extraction conditions that can lead to protein misfolding, aggregation, and loss of ligand binding activity. Over the past decade, cell-free (CF) expression of GPCRs has developed as a viable alternative to traditional cell-based methods, reducing the time taken to obtain purified GPCRs from months or weeks to days or hours (Yang *et al*., 2011; Rues *et al*., 2018; Köck *et al*., 2022). This has resulted in the publication of high-resolution cryo-EM structures of *E. coli* CF lysate-expressed full-length human β1-adrenoceptor (ADRB1) and histamine H2 receptor (H2R) in complex with G-proteins (Merino *et al*., 2025; Köck *et al*., 2024). In each case, GPCRs were co-translationally inserted into lipid nanodiscs that were present during CF expression to avoid the use of detergents. However, G-proteins were still produced using traditional insect cell approaches, thereby delaying the time to cryo-EM structure from the potential minimum offered by CF expression. In comparison to *E. coli* CF lysate, Protein synthesis Using Reconstituted Elements (PURE) reduces potential background contamination from host molecules, improves the ease of purification, and offers greater reproducibility, but has not been demonstrated to produce cryo-EM-ready GPCRs (Shimizu *et al*., 2001; Cui *et al*., 2022).

In this report, we combine ICL3-fusion construct design with Nuclera’s core *in vitro* transcription/translation PURE-based reagent and co-translational insertion into nanodiscs to rapidly accelerate the path to high-resolution cryo-EM GPCR structures. We present a 3.38 Å structure of PURE CF-expressed, antagonist-bound, human A_2A_R-β-lactamase fusion in complex with BLIP-II in nanodiscs, demonstrating for the first time the suitability of PURE-based CF expression to rapidly produce cryo-EM-ready GPCR complexes. Furthermore, by expanding this rapid expression and nanodisc screening approach to a wider set of 12 Class A-C GPCRs, we show that this detergent-free approach provides a generalizable pipeline for producing high-quality GPCRs for downstream biophysical and structural characterization.

## Results

### A_2A_R Purification and Nanodisc Screening on a Digital Microfluidic Device

Since Shah *et al*. demonstrated that insect cell-synthesized A_2A_R-β-lactamase in complex with BLIP-II can be readily resolved by cryo-EM, we adopted their construct strategy for testing in a PURE-based CF system (Shah *et al*., 2026). To rapidly assess whether A_2A_R can be co-translationally solubilized and purified, we screened expression and purification in the presence and absence of nanodiscs using the eProtein Discovery^TM^ digital microfluidics system (Nuclera). Linear eGene^TM^ DNA constructs encoding A_2A_R, A_2A_R-β-lactamase, and BLIP-II were designed for PURE-based CF expression using Nuclera CF Core reagent (CFCR). Constructs contained C-terminal split-GFP (DET) tag and Strep-tag II sequences for bimolecular fluorescence complementation detection (Romei and Boxer, 2019) and streptactin-based affinity purification, respectively.

Nanodiscs are phospholipid bilayers encased by engineered membrane scaffold proteins (MSPs) that solubilize and stabilize membrane proteins in detergent-free native-like lipid environments (Bayburt *et al*., 2002; Denisov *et al*., 2004). Nanodisc diameter is dictated by the length of the MSP belt used during assembly. GPCR insertion and folding is influenced by nanodisc lipid type and charge (Köck *et al*., 2022). Phosphatidylcholine (PC) lipids are common in eukaryotes and negatively charged phosphatidylglycerol (PG) lipids have been shown to increase translocon-independent membrane insertion, possibly through attraction of positive charges on the N-terminal part of the protein (Ridder *et al*., 2001). Cholesterol is important for the stabilization of certain GPCRs through direct protein-cholesterol interactions. Consequently, we screened three nanodisc lipid compositions (POPC/POPS, POPC/POPS/Cholesterol, and DOPG) and two MSP sizes (MSP1D1: ∼9.5 nm and MSP1E3D1: ∼13 nm).

Most GPCRs contain conserved extracellular disulfide bonds that are crucial for structural integrity and functional receptor folding. Because CF systems lack the endogenous oxidative folding machinery found within the endoplasmic reticulum of eukaryotic cells, protein disulfide isomerase (PDI) and oxidized glutathione (GSSG) were included in nanodisc-containing reactions to establish a redox-balanced environment that promotes correct disulfide bond formation. Expression and purification in the presence of nanodiscs and PDI + GSSG mix was compared against reactions containing CFCR and buffer alone. A condition with CFCR plus CaCl_2_ was included for BLIP-II, as it coordinates multiple Ca^2+^ ions.

As expected for a soluble protein, BLIP-II expressed efficiently and was largely unaffected by the presence or absence of nanodisc additives (Figure S1). A_2A_R and A_2A_R-β-lactamase displayed similar expression levels across all conditions, confirming that insertion of the β-lactamase fusion into ICL3 does not adversely affect protein synthesis in CFCR. The presence of nanodisc additives only moderately increased average expression yields compared to reactions performed in their absence (Figure 1a).

**Figure 1.**
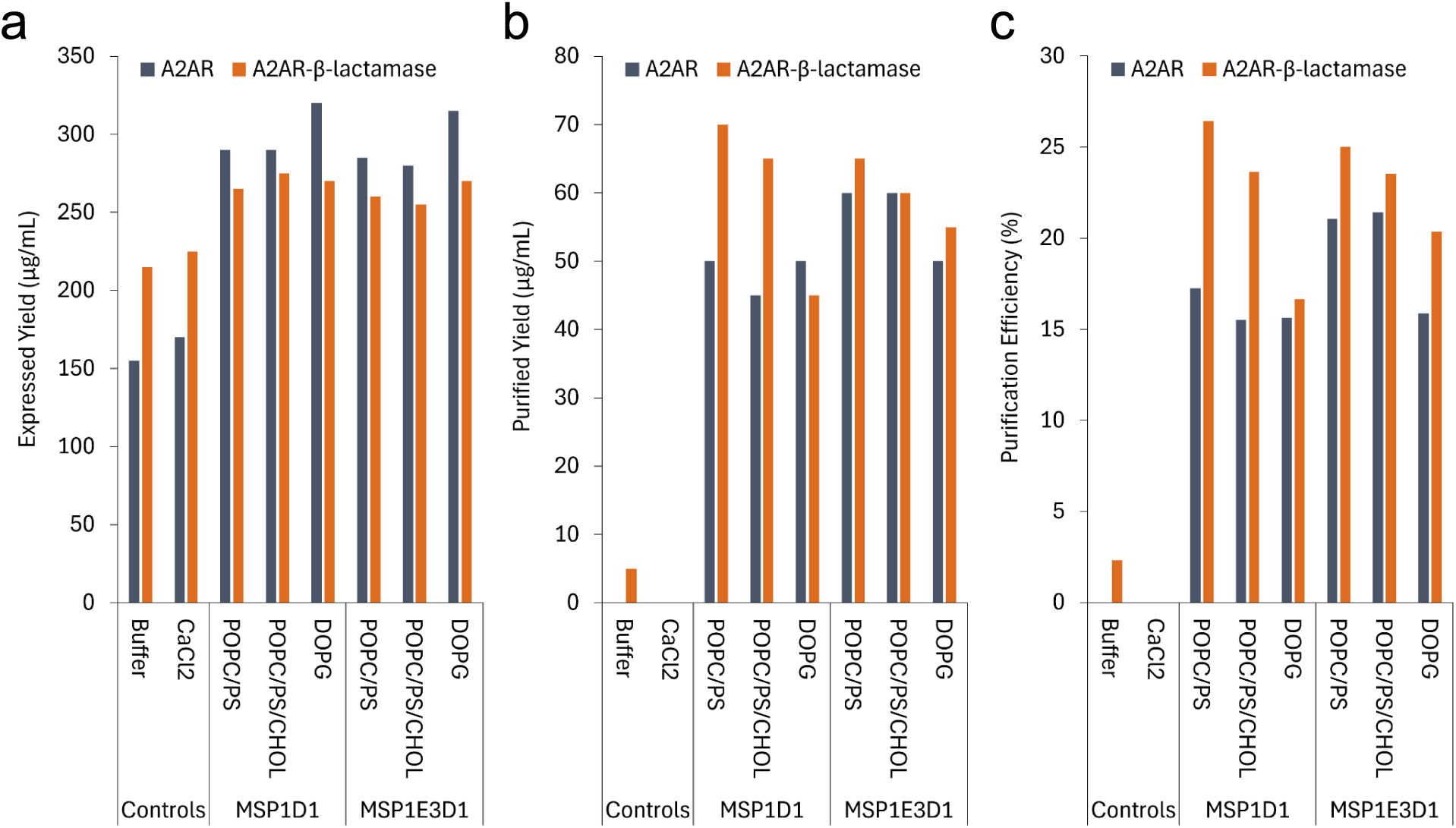
eProtein Discovery^TM^ screen of wild-type A_2A_R (gray) and A_2A_R-β-lactamase (orange) expression (**a**), purification (**b**), and purification efficiency (**C**) in the absence and presence of nanodiscs. Soluble control (BLIP-II) expression and purification data are provided in **Figure S1**. All nanodisc reactions contained PDI + GSSG mix.

Purified yields of BLIP-II were consistent across all tested conditions, reaching a final purified yield of 150 µg/mL with a purification efficiency of ∼30% (Figure S1). In contrast, A_2A_R and A_2A_R-β-lactamase expressed in the absence of nanodisc additives failed to purify to meaningful levels (Figure 1b). In the presence of nanodiscs and PDI + GSSG mix, purification yields for the wild-type and β-lactamase fusion protein were comparable, showing only minor variation across the different nanodisc types tested. Average purified yields across nanodisc conditions were 52.5 µg/mL for wild-type A_2A_R and 60.0 µg/mL for A_2A_R-β-lactamase (Figure 1b), corresponding to purification efficiencies of 18% and 23%, respectively (Figure 1c).

These results demonstrate that BLIP-II, wild-type A_2A_R, and A_2A_R-β-lactamase can be efficiently expressed and purified using CFCR. Successful affinity purification of A_2A_R constructs was strictly dependent on the presence of nanodiscs during translation, confirming effective co-translational stabilization of these membrane receptor complexes.

### A_2A_R-β-lactamase FSEC analysis and Complex Formation with BLIP-II

Previous studies using *E. coli* lysate-based CF expression demonstrated that size-exclusion chromatography (SEC) effectively resolves properly folded GPCR-nanodisc complexes from soluble aggregates and misfolded species that co-purify during affinity chromatography (Köck *et al*., 2022). To assess the biophysical quality and solution behaviour of A_2A_R, A_2A_R-β-lactamase, and BLIP-II purified from CFCR, expression reactions were scaled up in-tube for analytical fluorescence-detection SEC (FSEC) analysis. The eProtein Discovery™ screen identified several nanodisc formulations that supported GPCR expression, providing flexibility to select conditions based on downstream application requirements. MSP1E3D1 DOPG was selected for scale-up based on its expression performance and established precedent in enabling high-resolution cryo-EM structure determination of GPCRs produced using *E. coli* lysate-based CF systems (Merino *et al*., 2025; Köck *et al*., 2024).

A_2A_R, A_2A_R-β-lactamase, and BLIP-II were expressed independently and purified via Strep-tag II affinity capture prior to FSEC analysis. Purified yields were consistent with values measured on the eProtein Discovery^TM^ system (screen vs scale up; BLIPII: 0.16 vs 0.11 mg/mL; A_2A_R: 0.05 vs 0.07 mg/mL; A_2A_R-β-lactamase 0.06 vs 0.06 mg/mL) (Figure 2a). Both wild-type A_2A_R and A_2A_R-β-lactamase eluted earlier than empty nanodiscs and within the expected molecular weight range for monomeric GPCRs encapsulated in MSP1E3D1 DOPG nanodiscs (Figure 2 b and c and Figure S2a). As previously described for CF-expressed GPCRs (Köck *et al*., 2022), a proportion of aggregated material was detected, accounting for 35% and 36% of the total fluorescence for wild-type A_2A_R and A_2A_R-β-lactamase, respectively. BLIP-II eluted as a sharp monodisperse peak corresponding to the expected molecular weight of the soluble protein (Figure 2 b and c).

**Figure 2.**
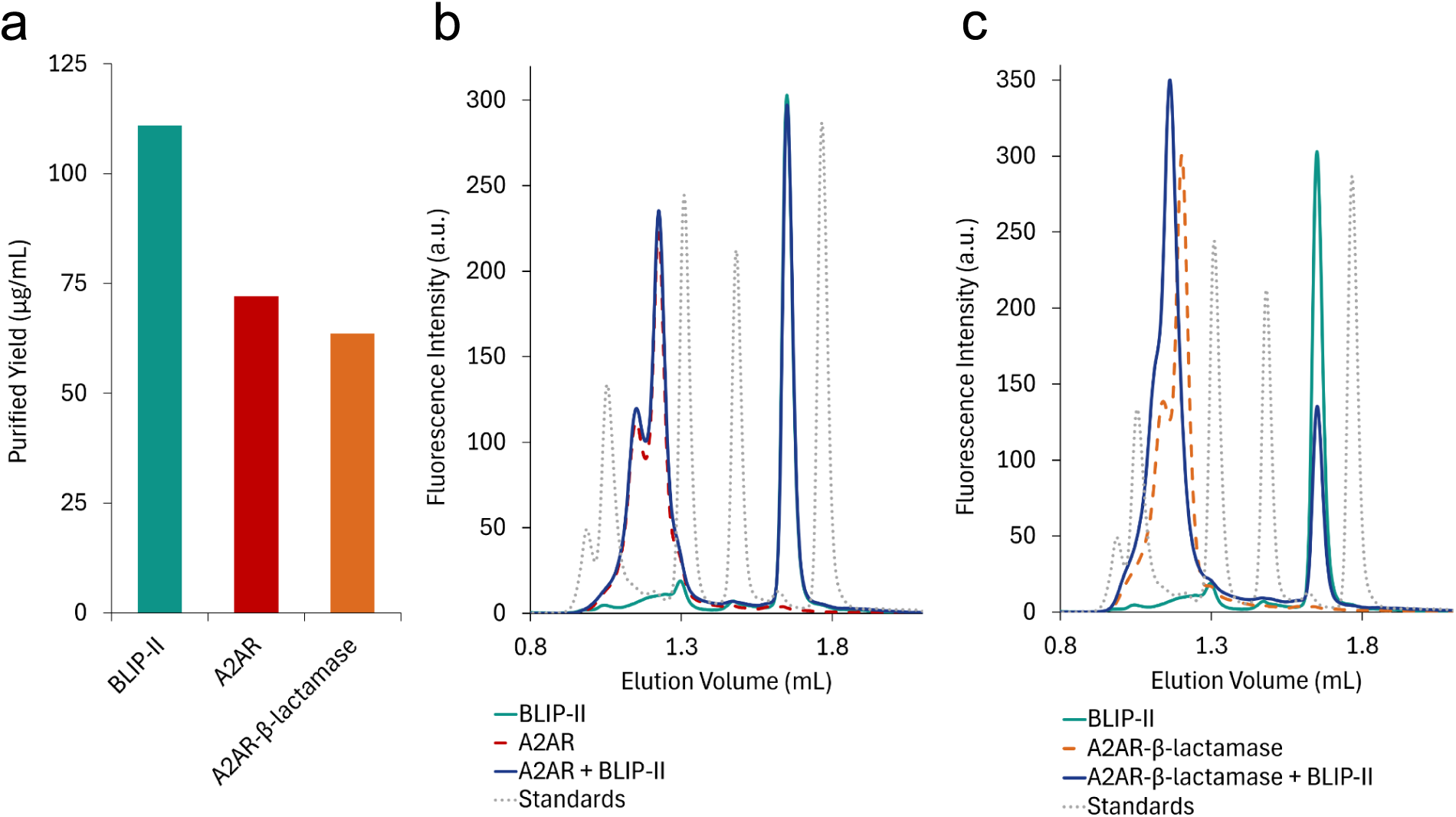
Scaled-up purification and FSEC confirmation of A_2A_R-β-lactamase-BLIP-II complex formation. (**a**) Purified yields of BLIP-II, A_2A_R, and A_2A_R-β-lactamase after strep-tactin purification. (**b**) FSEC traces for A_2A_R (red, dashed), BLIP-II (cyan), and A_2A_R with BLIP-II (blue). (**c**) FSEC traces for A_2A_R-β-lactamase (orange, dashed), BLIP-II (cyan), and A_2A_R-β-lactamase with BLIP-II (blue). Dotted gray traces indicate molecular weight standards (Waters, BEH200 SEC Test Mix) corresponding to 660 kDa, 150 kDa, 66.4 kDa, and 17 kDa.

To confirm functional complex assembly, A_2A_R-β-lactamase and BLIP-II were mixed prior to FSEC. Co-incubation resulted in a distinct retention time shift of the A_2A_R-β-lactamase peak towards a higher molecular weight, accompanied by an increase in peak height and a concomitant depletion of the free BLIP-II peak (Figure 2c). In contrast, mixing wild-type A_2A_R (lacking the ICL3 β-lactamase insertion) with BLIP-II produced no peak shift or change in peak intensity (Figure 2b). These results confirm that BLIP-II binds stably and specifically to the A_2A_R-β-lactamase fusion protein in nanodiscs.

### Structure Determination of A_2A_R-β-lactamase in Complex with BLIP-II in MSP1E3D1 DOPG Nanodiscs

To generate sufficient material for single-particle cryo-EM grid vitrification, A_2A_R-β-lactamase and BLIP-II expression was scaled up to 2 mL and 1 mL CFCR reactions, respectively. Following Strep-tag II affinity purification, the proteins were concentrated to ∼100 µL, yielding final concentrations of 1.25 mg/mL for A_2A_R-β-lactamase and 0.77 mg/mL for BLIP-II. Purified proteins were combined at a 1:1.2 molar ratio (8 µM A_2A_R-β-lactamase:9.6 µM BLIP-II) and supplemented with 15 µM ZM241385 antagonist to form the complex before plunge-freezing. Supplementation with ZM241385 did not impact the FSEC profiles of A_2A_R or A_2A_R-β-lactamase mixed with BLIP-II (Figure S2 b and c).

Single-particle cryo-EM analysis of the PURE-synthesized A_2A_R-β-lactamase–BLIP-II complex in MSP1E3D1 DOPG nanodiscs yielded a reconstruction at a global resolution of 3.38 Å from 47,986 particles (Figure 3a, Figure S3 and Table S3). The map allowed for confident docking of the high-resolution model PDB 9T9P with minimal further refinement. The map clearly resolves A_2A_R residues 8–146, 163–214 and 220–303, the complete β-lactamase insert, BLIP-II residues 41–311 and the bound antagonist ZM241385 (Figure 3c). Transmembrane helix 5 (TM5) continues without interruption into the N-terminal helix of β-lactamase, and the C-terminal β-lactamase helix extends directly into TM6, creating a rigid connection between the receptor and the fiducial module, as designed (Shah *et al*., 2026). Residues 147–162 of extracellular loop 2 (ECL2) and the C-terminal tail beyond residue 303 were not resolved, consistent with their flexibility. ECL2 was similarly poorly defined in the insect cell-derived structure (Shah *et al*., 2026), and retaining the full-length C-terminus in our construct did not prevent high-resolution reconstruction.

**Figure 3.**
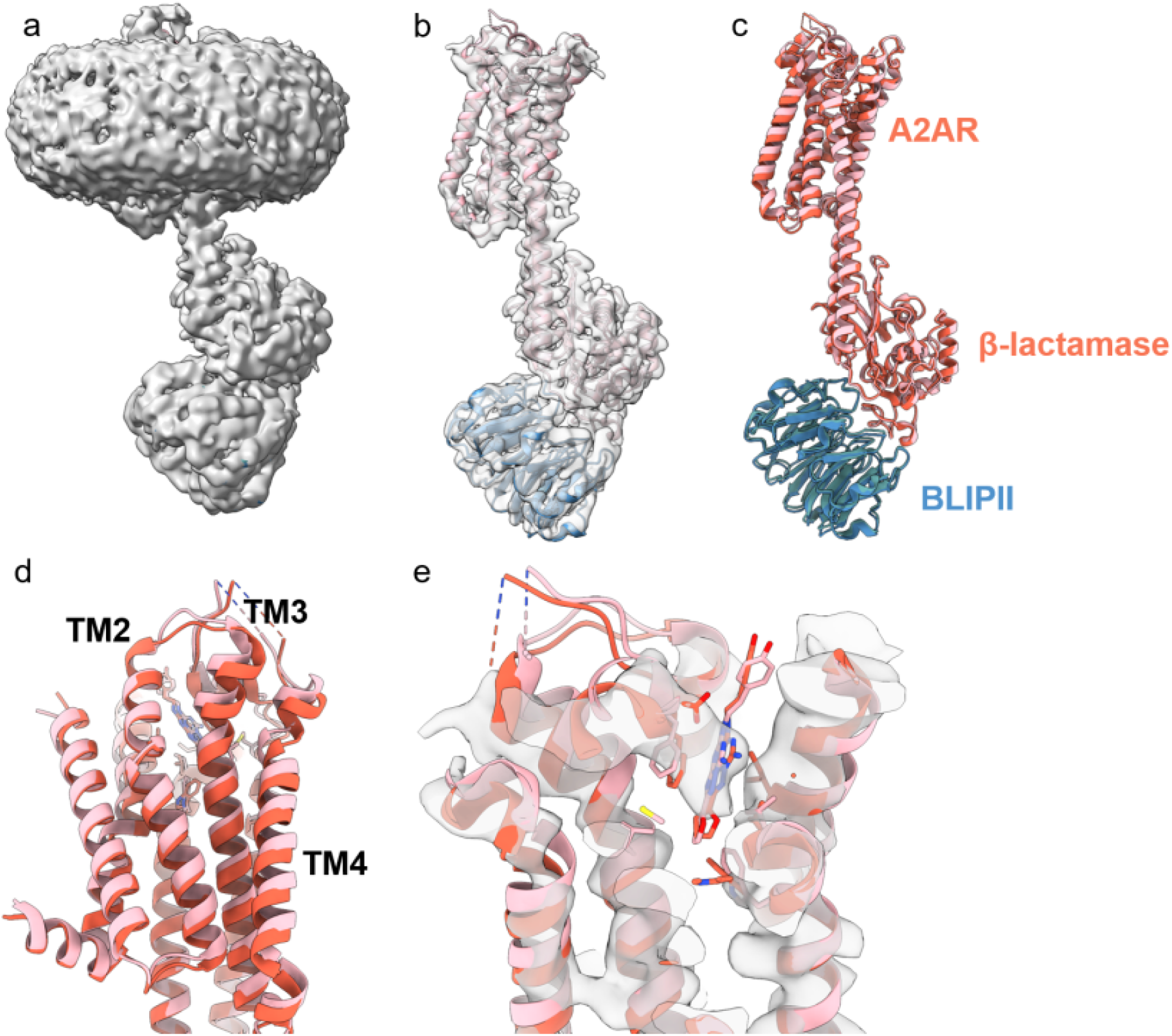
Structure of A2AR-β-lactamase in complex with BLIP-II in MSP1E3D1 DOPG Nanodiscs. All panels are shown in side view. (**a**) Cryo-EM map of A2AR–β-lactamase at 3.38 Å resolution. (**b**) Reference model (PDB 9T9P; Shah et al.) manually fitted into the map; nanodisc density was removed for clarity. (**c**) Superposition of the reference model (pink/steel blue) and the A2AR–β-lactamase model refined in this study (tomato/cadet blue). The models agree closely, with the main differences in TM2–TM4. (**d**) Close-up of TM2–TM4. (**e**) Close-up of the ligand fitted into the cryo-EM density.

The overall architecture closely matches that of the detergent-solubilized, insect cell-expressed complex (PDB 9T9P; Shah *et al*., 2026), with a Cα root-mean-square deviation (RMSD) of 0.99 Å over 803 residues and 0.85 Å over the 227 residues of the A_2A_R transmembrane helices (Figure 3c). Differences were confined to the extracellular ends of TM2 and TM3 around ECL1, to ECL2 and ECL3 (Cα deviations up to ∼2.3 Å), and to a small (∼3°) rigid-body tilt of the β-lactamase–BLIP-II module relative to the receptor, which displaces BLIP-II by ∼3.7 Å (Cα RMSD after superposition on the transmembrane helices) and may reflect the different membrane mimetics (Figure 3c and 3d). Map quality followed a gradient from the fiducial module towards the receptor (Figure 3a, 3b and Figure S3). β-lactamase and BLIP-II were the best-resolved components. Within the receptor, TM5 and TM6, which are directly continuous with the fusion helices, showed the strongest density with well-defined side chains, and TM2, TM3 and TM7 were well defined along their length. Density was weakest for the peripheral, lipid-facing TM1 and TM4, where many side chains were poorly resolved. The transmembrane bundle was encircled by a continuous, ∼4 nm-thick belt of unmodelled density corresponding to the DOPG bilayer and MSP1E3D1 scaffold, with the receptor positioned centrally in the disc. Notably, comparable overall resolution was achieved with approximately 12-fold fewer particles than were used for 9T9P (47,986 versus 597,000 particles, yielding 3.38 Å versus 3.22 Å consensus resolution; Shah *et al*., 2026).

Obvious, but poorly defined non-protein density was present in the orthosteric binding pocket, into which ZM241385 was docked (Figure 3e). Density for the triazolotriazine core and furan ring was comparable to that of the surrounding side chains, whereas the 4-hydroxyphenylethylamino tail, which projects towards the solvent-exposed extracellular vestibule, was weaker. This mirrors observations for 9T9P, in which the bicyclic core and furan ring could be placed confidently only after focused refinement of the receptor (Shah *et al*., 2026). The antagonist adopts the canonical binding mode: the bicyclic core is sandwiched between Phe168^ECL2^ and Leu249^6.51^, the furan ring extends deep into the pocket towards Trp246^6.48^ and His250^6.52^, and the exocyclic amine faces Glu169^ECL2^ and Asn253^6.55^. After superposition on the transmembrane helices, ZM241385 occupies the same site as in 9T9P (heavy-atom RMSD 1.7 Å), and the 9T9P ligand pose fits our map equally well, indicating that the binding modes are indistinguishable at the current resolution. Side-chain differences among the 15 residues lining the pocket are likewise within the precision expected at this resolution. Together, these data demonstrate that A_2A_R synthesized in the PURE-based system and co-translationally inserted into nanodiscs adopts a native fold and binds ZM241385 in the same manner as receptor produced in insect cells.

### Broad Application of PURE Co-Translational Nanodisc Insertion to Diverse GPCR Classes and Orphan Targets

To evaluate the generalizability of the PURE CF co-translational nanodisc insertion platform beyond the A_2A_R benchmark, we expanded our screening panel to cover additional examples and structural classes within the GPCR superfamily. This expanded set included non-orphan Class A receptors (C-C chemokine receptor 10, CCR10; C-X-C chemokine receptor 1, CXCR1; β_2_-adrenergic receptor, ADRB2; and cannabinoid receptor 2, CB2) alongside representative targets from Class B, (glucagon receptor, GCGR), and Class C (retinoic acid-induced protein 3, RAI3).

Beyond characterized targets, uncharacterized orphan GPCRs represent high-value targets for this detergent-free co-translational stabilization system. Because their natural ligands are unknown, orphan GPCRs cannot be thermostabilized using conventional small-molecule orthosteric ligands or stabilized via signalling protein complexes cell-based extraction. Having established that our workflow yields functionally intact, ligand-responsive GPCRs with our antagonist-bound A_2A_R-β-lactamase structure, we tested whether high-quality unliganded (apo) orphan GPCRs could be generated by incorporating six Class A orphan receptors into our panel: GPR61, GPR85, GPR88, GPR146, GPR151, and GPR174.

Because extracellular domain architecture and N-terminal folding can influence translation efficiency and membrane incorporation, we evaluated three N-terminal construct variants for each target: (1) native untagged, (2) an N-terminal BRIL fusion tag to enhance soluble folding, and (3) an N-terminal fusion with the amphipathic membrane-anchoring helix of dihydroorotate dehydrogenase (DHODH) to facilitate initial nanodisc association. All construct variants were rapidly screened across distinct nanodisc compositions on the eProtein Discovery™ to identify optimal expression conditions.

Reaction conditions producing the highest yield of purified protein for each target (Table S1) were scaled up in-tube, and analytical FSEC was used to assess protein quality and aggregation levels. All 12 receptors were successfully expressed, co-translationally inserted into nanodiscs, and purified in the unliganded (apo) state. Purification yields for all targets were similar to or greater than those obtained for A_2A_R (Figures 2 and Figure S4). The non-orphan panel comprising Class A (Figure 4a-d), B (Figure 4e), and C (Figure 4f) examples eluted as distinct, well-defined peaks corresponding to monomeric GPCR-nanodisc complexes. Aggregation levels varied, but remained low across all three classes. Similarly, six orphan GPCRs exhibited sharp, symmetrical elution profiles with high fluorescence intensities in the expected molecular weight range for single receptor-nanodisc assemblies (Figure 4g–l). Notably, GPR61 (Figure 4g), GPR146 (Figure 4j) and GPR151 (Figure 4k) yielded near-monodisperse peaks with negligible high-molecular-weight aggregate species.

**Figure 4.**
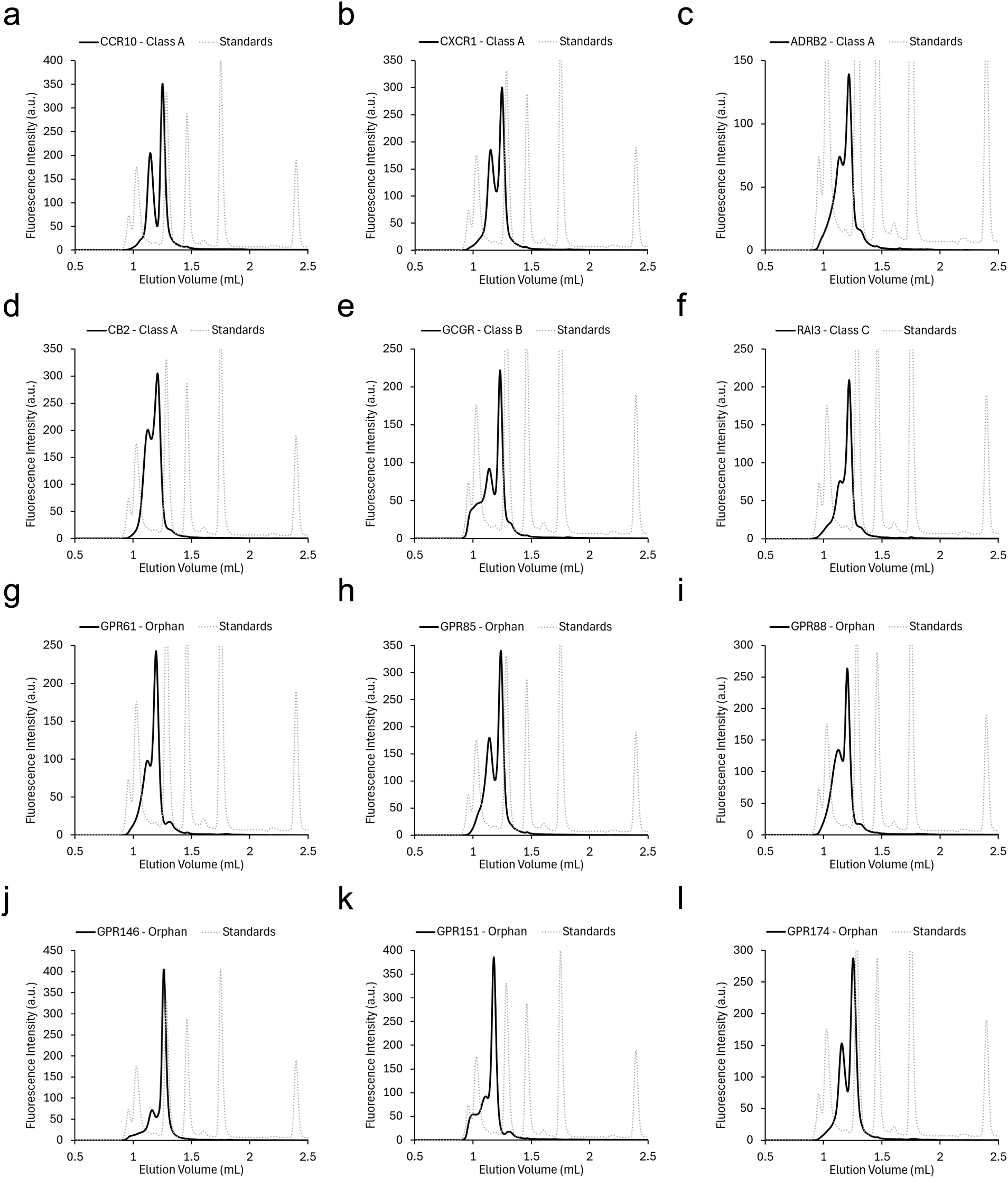
Analytical FSEC analysis of cell-free expressed and affinity-purified GPCRs in nanodiscs. Tested constructs span Class A receptors including **CCR10** (**a**), **CXCR1** (**b**), **ADRB2** (**c**), and **CB2** (**d**); Class B receptor **GCGR** (**e**); Class C receptor **RAI3** (GPRC5A) (**f**); as well as orphan GPCRs **GPR61** (**g**), **GPR85** (**h**), **GPR88** (**i**), **GPR146** (**j**), **GPR151** (**k**), and **GPR174** (**l**). Reconstructed receptor–nanodisc assemblies are plotted in solid black traces. Dotted gray traces show molecular weight standards (Waters, BEH200 SEC Test Mix) corresponding to 660 kDa, 150 kDa, 66.4 kDa, 17 kDa, and 112 Da.

Across the entire 12-receptor panel, the predominant peak consistently matched the expected molecular weight profile for single receptor-nanodisc assemblies, with minimal high-molecular-weight aggregation. The biophysical quality of the expanded GPCR panel was comparable to, or exceeded, that of A_2A_R-β-lactamase, which proved sufficient for high-resolution structure determination. These findings demonstrate that PURE-driven co-translational nanodisc insertion is a robust, target-agnostic platform capable of generating detergent-free, structural-grade preparations across diverse GPCR structural families and unliganded orphan targets.

## Discussion

High-resolution single-particle cryo-EM has transformed structural biology (Cheng, 2015), yet antagonist-bound and unliganded GPCRs remain challenging targets (Zhang *et al*., 2022). Because GPCRs feature small, membrane-embedded domains with minimal extracellular mass, achieving reliable particle alignment typically requires coupling to heterotrimeric G-proteins and/or stabilizing antibody fragments (Zhang *et al*., 2022). While engineering rigid fiducials can bypass the requirement for G-protein coupling (Singh *et al*., 2026; Shah *et al*., 2026; Collu *et al*., 2026), upstream protein production remains a central bottleneck. Traditional insect or mammalian cell-based expression pipelines can require weeks of construct optimization and rely on detergent solubilization, which can cause protein misfolding, loss of allosteric regulation, and structural instability in the absence of high-affinity ligands (Milic and Veprintsev, 2015; Tate, 2001).

In contrast to cell-based approaches, CF expression of GPCRs permits co-translational stabilization with lipid nanodiscs, completely avoiding detergent exposure and preserving receptor stability (Yang *et al*., 2011; Rues *et al*., 2018; Köck *et al*., 2022). The recent cryo-EM structures of *E. coli*-lysate-produced GPCRs from the Bernhard laboratory represent significant advances in establishing CF platforms for membrane protein structural biology (Köck *et al*., 2024; Merino *et al*., 2025). However, those structures relied on insect-cell-synthesized trimeric G-proteins, which consequently slowed protein production timelines compared to an entirely CF pipeline (Köck *et al*., 2024; Merino *et al*., 2025). Moreover, these strategies are not applicable to apo or antagonist-bound receptor states, which remain significantly underrepresented across available cryo-EM structures due to their intrinsic structural flexibility and lack of stabilizing binding partners (Singh *et al*., 2026). Determining apo GPCR structures is critical for drug discovery; unliganded conformations reveal basal receptor dynamics, uncover self-inhibitory or tethered-agonist mechanisms (Lin *et al*., 2020), and expose transient allosteric binding pockets obscured in ligand-bound active states (Hollingsworth *et al*., 2019). Resolving this technical gap is essential for expanding the structural landscape of uncharacterized orphan receptors and driving rational antagonist or allosteric drug development (Mao *et al*., 2024).

In this study, we combined recent advances in rigid fiducial design (Shah *et al*., 2026) with PURE-based co-translational GPCR stabilization to determine a 3.38 Å structure of antagonist-bound A_2A_R-β-lactamase in complex with BLIP-II in nanodiscs using a completely CF workflow. By utilizing automated, nanoliter-scale construct and nanodisc screening on the eProtein Discovery^TM^ digital microfluidic system, coupled with small-scale in-tube expression, purification, and FSEC analysis (Kawate and Gouaux, 2006), we rapidly validated cryo-EM readiness of A_2A_R-β-lactamase. Crucially, microfluidic screening confirmed that co-translational insertion into nanodiscs was strictly required for A_2A_R solubilization and affinity purification, de-risking reaction scale-up before committing higher reagent volumes. Moreover, analytical FSEC analysis of small-scale scale-up reactions confirmed the suitability of the PURE-based CF system for producing high-quality monomeric GPCR-nanodisc complexes. These validation steps allowed us to calculate the required scale-up volumes based on purified yields and biophysical quality. Subsequent 1-2 mL expression reactions of A_2A_R-β-lactamase and BLIP-II yielded sufficient material for high-resolution cryo-EM structure determination directly from a single affinity-purification step. Unlike protein produced in insect cells, CF-synthesized A_2A_R-β-lactamase did not require truncation of the C-terminus to obtain sufficient quantities for cryo-EM. In total, the process from initial construct generation and microfluidic screening on the eProtein Discovery^TM^ system through scale up and FSEC analysis to cryo-EM grid vitrification took four days, representing a dramatic acceleration compared to traditional cell-based production pipelines.

The 3.38 Å structure of A_2A_R-β-lactamase–BLIP-II shows that GPCRs synthesized in the PURE system and co-translationally inserted into nanodiscs are structurally equivalent to receptors produced by conventional cell-based methods. The receptor adopts the same fold as the detergent-solubilized, insect cell-expressed complex (Shah *et al*., 2026), with sub-ångström deviations across the transmembrane helices, and ZM241385 occupies the orthosteric pocket in the canonical binding mode, confirming that CF-synthesized A_2A_R is correctly folded and competent for ligand binding. Unlike previous CF-derived GPCR structures, which required insect cell-produced G-proteins and captured agonist-bound active states (Köck *et al*., 2024; Merino *et al*., 2025), every component of this complex was produced cell-free, and the structure represents an antagonist-bound state that is inaccessible to G-protein-based approaches. Notably, a resolution comparable to that of 9T9P was reached from approximately 12-fold fewer particles, despite retention of the full-length, disordered C-terminus. Although differences in data collection and processing preclude a direct comparison, this suggests that nanodisc-embedded, CF-synthesized material is sufficiently homogeneous for efficient cryo-EM analysis. The lipid bilayer also provides a more native environment than detergent micelles, and the small difference in the orientation of the fiducial module relative to the receptor may reflect this. Indeed, recent NMR conformational profiling of A_2A_R demonstrated that conventional detergent environments can significantly distort receptor conformational equilibria compared to nanodiscs, which maintain native-like structural dynamics (Mendoza-Hoffmann *et al*., 2024).

Map quality was highest for the fiducial module and decreased towards the receptor, with the peripheral TM1 and TM4 and the extracellular loops least well defined, and density for ZM241385, particularly its phenylethylamino tail, was weaker than for the protein core. A similar gradient was observed for 9T9P, in which focused refinement of the receptor was required to place the ligand confidently (Shah *et al*., 2026), and probably reflects residual flexibility between the receptor and the fusion together with the contribution of the surrounding lipid and scaffold density to particle alignment. At the current resolution, the ligand pose is consistent with previous structures, but detailed interactions such as hydrogen-bonding networks and ordered water molecules, which are important for structure-based drug design, cannot be resolved. Because the CF workflow is readily scalable, larger datasets combined with focused refinement of the receptor, further rigidification of the fusion junctions, or the addition of binders that increase the mass of the fiducial module are expected to improve receptor and ligand density. Nonetheless, the present structure establishes that the complete path from linear DNA template to a cryo-EM structure of an antagonist-bound GPCR complex can be achieved using CF synthesis alone.

Beyond validating this workflow with A_2A_R-β-lactamase, we demonstrated its broader applicability by evaluating an expanded panel of 12 additional receptors, covering Class A, B, and C targets as well as orphan GPCRs. Crucially, FSEC analysis of this 12-target panel revealed protein quality similar to A_2A_R-β-lactamase, which proved sufficient for high-resolution cryo-EM structure determination. These data demonstrate that direct scale-up of top-performing eProtein Discovery^TM^ screen conditions reliably yields structural-grade GPCR-nanodisc complexes following single-step affinity purification. Moreover, because these receptors remain stable in the unliganded state, they provide versatile substrates for both apo and ligand-bound structural and biophysical characterization.

While the present study utilized the A_2A_R-β-lactamase construct as a structural benchmark to establish proof-of-concept, the rigid fiducial fusion strategy could be extended to the broader target panel to facilitate downstream single-particle cryo-EM alignment of both antagonist-bound and apo receptor conformations (Shah *et al*., 2026; Singh *et al*., 2026). Inserting fiducial domains into intracellular loops could influence protein folding, membrane insertion efficiency, or ligand binding activity (Chun *et al*., 2012). Shah *et al*. demonstrated that ICL3-β-lactamase fusion did not alter ligand binding of A_2A_R (Shah *et al*., 2026). When expanding this approach to other GPCR targets, any impact on ligand binding activity should be derisked and β-lactamase insertion junctions should be determined through target-specific optimization strategies. Combining AI construct-design with high-throughput microfluidic screening of engineered fiducial variants and detergent-free PURE-based co-translational insertion provides a scalable, generalizable framework to streamline the structure determination of challenging GPCRs. Ultimately, this approach provides a rapid pathway amenable to Design–Build–Test–Learn (DBTL) workflows, expanding the accessible membrane protein landscape to drive fundamental mechanistic discovery and accelerate therapeutic development across previously intractable GPCR targets.

## Materials and Methods

### Construction of eGene^TM^ Constructs

Genes of interest were retrieved from UniProt (Table S2). The sequence for A_2A_R-β-lactamase was adapted from Shah *et al*. (2026) without the C-terminal truncation (Shah *et al*., 2026). Linear DNA fragments encoding the genes were designed using eProtein Discovery^TM^ software (Nuclera) and purchased as standard gBlocks^TM^ or HiFi gene fragments (Integrated DNA Technologies) featuring 5′ 3C and 3′ TEV cleavage sequences (Table S2). eGene^TM^ constructs were assembled via PCR using the Linear eGene^TM^ Prep Kit Flexivariant^TM^ (Nuclera, Cat# NC3008) and Platinum^TM^ SuperFi II PCR MasterMix (Thermo Fisher Scientific, Cat# 12368010) according to the manufacturers protocol to append 5′ flanking elements (promoter, ribosome binding site, translation enhancer) and 3′ flanking elements (DET tag, Strep-tag II, and translation/transcription terminators). Assembled linear eGene^TM^ DNA was purified using AMPure XP DNA cleanup beads (Beckman Coulter, Cat#A63881) and quantified using Quant-iT^TM^ PicoGreen^TM^ assay kits (Thermo Fisher Scientific, Cat# P7581).

### eProtein Discovery^TM^ Screen

Multiplex cell-free protein synthesis (CFPS) screening was performed on an eProtein Discovery™ system (Nuclera) using eProtein Discovery™ Cartridges (Nuclera, Cat# NC3006). Cell-free blends were prepared using the Cartridge Reagent Kit: Membrane Protein (Nuclera, Cat# NC3013) supplemented with Additive Buffer, CaCl_2_, PDI + GSSG Mix, or nanodiscs (Nuclera eProtein Discovery™ GPCR Nanodisc Panel Screen Kit, Cat# NC3023). Screens were performed following manufacturer’s protocols and reagents. Protein expression and purifiability were automatically quantified on-cartridge using split-GFP fluorescence complementation and streptactin-bead-based purification following the manufacturer’s Membrane Protein Workflow.

### Scale up and FSEC Analysis

For FSEC analysis, GPCR expression was scaled up to 100 µL in reactions containing 60 µL Scale-Up Core (Nuclera, Cat# NC3011-2), 7.5 µL PDI + GSSG Mix (Nuclera, Cat# NC3005), 7.5 µL MSP1E3D1 DOPG (Nuclera, Cat# NC30128), and 25 µL linear eGene^TM^ DNA. For BLIP-II expression, PDI + GSSG Mix and nanodiscs were replaced with 15 µL CaCl_2_ additive (Nuclera, Cat# NC3005). Final concentrations of nanodiscs and eGene^TM^ DNA in the reaction were 75 µM and 5 nM, respectively.

Reactions were incubated at 29°C overnight for expression prior to supplementation with 0.2 mg/mL Monarch® RNase A (NEB, Cat# T3018L) and incubation for an additional 30 min. Expressed proteins were purified via Strep-tag II magnetic bead capture using Strep Beads (Nuclera, NC3011-1). Proteins were eluted in 100 µL of Elution Buffer (Nuclera, NC3011-2). Protein concentrations were quantified by split-GFP fluorescence complementation using the eProtein^TM^ Quantification Kit (Nuclera, Cat# NC3014) according to the manufacturer’s instructions.

FSEC was carried out at 25°C using a Waters ACQUITY Premier UPLC System equipped with an ACQUITY Premier Protein SEC Column (250 Å, 1.7 µm, 4.6 x 150 mm; Waters, 186009963). The column was equilibrated in filter-sterilized running buffer (20 mM sodium phosphate dibasic, 250 mM sodium chloride, 100 mM arginine, pH 7.5). Purified protein samples were filtered through a 0.22 µm membrane. A 10 µL volume of diluted A_2A_R (0.45 µM), A_2A_R-β-lactamase (0.45 µM), or BLIP-II (0.55 µM) was injected at a flow rate of 0.25 mL/min. Tryptophan fluorescence (Ex280/Em350 nm) was monitored over a 15 min run time. For the expanded GPCR panel, 2.5 µL of the purified protein was injected at a flow rate of 0.28 mL/min and tryptophan fluorescence (Ex280/Em350 nm) was monitored over a 12 min run time.

### Cryo-EM Sample Preparation

To prepare proteins for cryo-EM, A_2A_R-β-lactamase expression was scaled up to a 2 mL reaction containing 1200 µL Scale-Up Core (Nuclera, Cat# NC3011-2), 150 µL PDI + GSSG Mix (Nuclera, Cat# NC3005), 150 µL MSP1E3D1 DOPG (Nuclera, Cat# NC30128), and 500 µL linear eGene^TM^ DNA. BLIP-II expression was scaled up to a 1 mL reaction containing 600 µL Scale-Up Core (Nuclera, Cat# NC3011-2), 150 µL CaCl_2_ additive (Nuclera, Cat# NC3005), and 250 µL linear eGene^TM^ DNA.

Reactions were incubated at 29°C overnight and subsequently treated with 0.2 mg/mL Monarch® RNase A (NEB, Cat# T3018L) at 29°C for 30 min. Expressed proteins were purified via Strep-tag II magnetic bead capture using Strep Beads (Nuclera, NC3011-1) and eluted in 2 mL (A_2A_R-β-lactamase) or 1 mL (BLIP-II) of detergent-free Elution Buffer. Eluted proteins were concentrated to ∼100 µL using Amicon® Ultra 0.5 mL Centrifugal Filters with a 30 kDa MWCO (A_2A_R-β-lactamase; Merck Millipore, Cat# UFC503024) or 10 kDa MWCO (BLIP-II; Merck Millipore, Cat# UFC501024). Protein concentrations were quantified using the eProtein^TM^ Quantification Kit (Nuclera, Cat# NC3014) according to the manufacturer’s instructions.

To assemble the complex for Cryo-EM, purified A_2A_R-β-lactamase and BLIP-II were mixed at final concentrations of 8 µM and 9.6 µM, respectively, supplemented with 15 µM ZM241385 (Merck Millipore, Cat# Z0153) antagonist, and incubated on ice overnight prior to grid vitrification. For cryo-EM analysis of freeze-thawed material, aliquots stored at −80 °C were thawed on ice before grid preparation.

### Cryo-EM Grid Vitrification

A 3 µL aliquot of sample was applied onto Quantifoil® R1.2/1.3 Cu 300-mesh grids. Prior to sample application, grids were glow-discharged on both sides at 25 mA for 60 s per side. Grids were vitrified using a Vitrobot (ThermoFisher Scientific) operated at 4°C and 95% relative humidity, with a blotting time of 3 s and a blot force of -5.

### Cryo-EM Data Collection

Cryo-EM data were acquired on a Titan Krios transmission electron microscope (ThermoFisher Scientific) operated at 300 kV and equipped with a Selectris X energy filter and a Falcon 4i direct electron detector operated in electron counting mode. Images were recorded at a nominal magnification of 165,000×, corresponding to a calibrated pixel size of 0.729 Å pixel^−^¹. A 10 eV energy-filter slit width was used. A total of 31,799 movies were acquired using aberration-free image shift (AFIS) in EPU (ThermoFisher Scientific). Data were collected over a nominal defocus range of −1.8 to −0.6 μm in 0.2 μm increments.

### Cryo-EM image processing

Movies were pre-processed in real time using WARP (Tegunov and Cramer, 2019). Gain correction, motion correction, and contrast transfer function (CTF) estimation were performed prior to automated particle picking.

Particles were automatically picked using BoxNet2 and extracted in 384-pixel boxes at 0.729 Å pixel^−^¹. A total of 5,477,692 particles were picked and imported in cryoSPARC (v 5.0.6) (Punjani A. *et al*., 2017) for subsequent steps of data processing.

Two-dimensional classification was used to remove erroneous picks and identify classes corresponding to the A_2A_R–β-lactamase-BLIP-II complex. Subsequently, iterative rounds of three-dimensional classification and refinement identified a subset of 47,986 particles, which produced the highest-resolution reconstruction at an overall resolution of 3.38 Å. The resulting 3D density map aligned well with the structural architecture of the reference A_2A_R–β-lactamase model (PDB 9T9P). The PDB file 9T9P was manually fitted in ChimeraX (Pettersen EF *et al.,* 2021), and adjusted in Coot 0.9.8.95 (Emsley P. *et al*., 2010), and subsequently refined in Phenix 1.21 (Adams PD. *et al.,* 2010).

## Data availability

All biochemical, microfluidic screening, and size-exclusion chromatography datasets supporting the findings of this study are included within the main text and Supplementary Information. Cryo-EM atomic coordinates and atomic density maps for the reported structures are in preparation for deposition to the Protein Data Bank (PDB) and will be made publicly available upon peer-reviewed publication. Accession codes will be provided prior to formal publication.

## Acknowledgements

This study was funded by Nuclera Ltd. We acknowledge the School of Biological Sciences Cryo-Electron Microscopy Facility (SBS Cryo-EM Facility), where all cryo-EM experiments were performed. The facility’s instrumentation was funded by the Wellcome Trust (grant numbers 206171/Z/17/Z and 202905/Z/16/Z) and the University of Cambridge.

## Author Contributions

TAG conceived the study, developed the workflow, performed scale up experiments, FSEC analysis, wrote the manuscript, and prepared figures. AJK and ZBG performed eProtein Discovery^TM^ screen experiments, assisted in workflow development, and performed data analysis. MMM performed eProtein Discovery^TM^ screen experiments, assisted in FSEC analysis experiments, workflow and strategy development. GP, SWH, LC, and DYC performed cryo-EM analysis, prepared figures, and assisted in manuscript preparation. TWBO provided technical strategy guidance and support.

