## Supplementary Information for "Rapid Production and Cryo-EM Structure Determination of GPCR-Nanodisc Complexes via Cell-Free Synthesis"

| Name | Class | Construct | Additive 1 | Additive 2 |
|---|---|---|---|---|
| CCR10 | A | NoTag | MSP1E3D1 DOPG | PDI + GSSG Mix |
| CXCR1 | A | NoTag | MSP1E3D1 DOPG | PDI + GSSG Mix |
| ADRB2 | A | NoTag | MSP1E3D1 DOPG | PDI + GSSG Mix |
| CB2 | A | BRIL | MSP1E3D1 DOPG | PDI + GSSG Mix |
| GCGR | B | NoTag | MSP1E3D1 DOPG | PDI + GSSG Mix |
| RAI3 | C | NoTag | MSP1E3D1 DOPG | PDI + GSSG Mix |
| GPR61 | Orphan A | NoTag | MSP1E3D1 DOPG | PDI + GSSG Mix |
| GPR85 | Orphan A | BRIL | MSP1E3D1 DOPG | PDI + GSSG Mix |
| GPR88 | Orphan A | BRIL | MSP1E3D1 DOPG | PDI + GSSG Mix |
| GPR146 | Orphan A | NoTag | MSP1E3D1 DOPG | PDI + GSSG Mix |
| GPR151 | Orphan A | DHODH | MSP1E3D1 DOPG | PDI + GSSG Mix |
| GPR174 | Orphan A | NoTag | MSP1E3D1 DOPG | PDI + GSSG Mix |

***Table S1.*** *Conditions selected from the eProtein Discovery™ screen for scale-up expression, purification, and FSEC analysis (Figure 4).*

| Name | Source | Modifications | gBlock Sequence |
|---|---|---|---|
| A2AR | P2974 | N-term Met removed | CTCGAGGTTCTGTTCCAAGGACCTCCGATCATGGGGTCGTCCGTATATATCACGGTAGAACTGGC<br>TATTGCGGTCTTGGCTATCTTAGGAAACGTCTTGGTTTGCTGGGCGGTGTGGTTAAACTCGAACT<br>TGCAAAACGTAACTAATTACTTCGTGGTTTCCTTGGCCGCTGCAGACATTGCAGTCGGGGTCTTG<br>GCTATTCCCTTCGCTATCACAATATCTACTGGTTTTTGTGCGGCTTGTCATGGCTGCTTGTTTAT<br>CGCATGTTTTGTTCTTGTATTGACACAAAGCTCTATTTTTAGTCTGCTTGCAATTGCCATCGACC<br>GTTACATCGCCATCCGGATTCCCTTAAGATACAACGGCCTGGTGACTGGGACGCGCGCTAAAGGA<br>ATTATAGCAATCTGTTGGGTCTTATCGTTTGCGATAGGGCTTACGCCAATGCTGGGGTGGAACAA<br>TTGCGGACAACCTAAGGAGGGTAAAAACCACTCCCAGGGGTGTGGTGAGGGTCAAGTGGCTTGTC<br>TGTTTGAAGATGTAGTCCCTATGAATTATATGGTTTACTTTAACTTCTTCGCGTGCGTGCTTGTC<br>CCCCTGTTACTTATGTTAGGTGTTTACCTGCGGATATTTCTTGCCGCCCGGCGTCAATTAAAGCA<br>AATGGAATCGCAGCCACTGCCGGGCGAACGGGCCCGGTCTACCCTTCAGAAAGAGGTACATGCCG<br>CCAAATCGTTGGCAATCATCGTAGGACTTTTTGCTTTATGTTGGTTGCCTTTGCATATAATCAAC<br>TGTTTTACGTTTTTTTGCCCAGATTGCAGTCACGCCCCATTGTGGCTGATGTACCTTGCCATAGT<br>TCTGTCGCACACCAACTCTGTAGTTAACCCGTTCATATATGCGTATAGAATTAGAGAATTCCGGC<br>AAACCTTCCGCAAAATAATTCGCTCACATGTTCTTAGACAGCAAGAACCCTTTAAAGCCGCTGGG<br>ACAAGTGCTCGCGTGTTGGCGGCCCACGGTTCAGATGGAGAGCAAGTATCTTTACGGTTGAATGG<br>ACACCCGCCAGGTGTGTGGGCCAACGGTTCTGCTCCCCACCCTGAAAGACGGCCTAACGGTTACG<br>CATTAGGATTGGTGTCAGGTGGGTCGGCGCAAGAGTCCCAAGGGAACACCGGACTGCCGGACGTA<br>GAATTACTTTCGCACGAATTAAAGGGCGTTTGTCCCGAACCCCCCGGCCTGGATGATCCACTGGC<br>GCAAGATGGTGCCGGAGTATCAGAGAACCTGTACTTCCAGAGC |
| A2AR-β-lactamase | Shah et al., 2026 | No C-term truncation | CTCGAGGTTCTGTTCCAAGGACCTCCTATTATGGGAAGTTCGGTTTATATCACCGTAGAATTGGC<br>AATCGCGGTTTTGGCTATTTTAGGAAACGTGCTGGTATGCTGGGCCGTGTGGCTTAATAGCAATT<br>TGCAGAATGTCACGAATTATTTCGTTGTGTCTCTGGCGGCAGCTGATATAGCTGTAGGTGTATTA<br>GCAATCCCATTTGCTATAACAATCTCAACAGGGTTCTGTGCAGCATGTCATGGGTGTCTTTTCAT<br>TGCTTGCTTCGTTTTAGTACTGACTCAGAGCAGTATCTTTTCATTGCTTGCGATTGCCATAGACA<br>GATATATTGCGATACGTATCCCCTTGCGGTATAACGGTTTGGTCACAGGAACACGCGCCAAGGGG |

| | | | |
|---|---|---|---|
| | | | ATAATAGCTATATGCTGGGTGCTGTCTTTTGCTATTGGGCTTACGCCAATGTTGGGATGGAATAA<br>TTGCGGACAGCCAAAAGAAGGTAAGAATCATTCGCAAGGATGTGGAGAGGGCCAGGTCGCGTGTT<br>TGTTTGAGGATGTTGTGCCCATGAACTACATGGTGTATTTTAATTTCTTTGCATGCGTTTTAGTT<br>CCTCTGTTGCTGATGCTGGGGGTGTATTTGAGAATATTCCTTGCAGCGCGTCGTCAGTTGAAACA<br>AATGGAGTCTCAAGCCAAGGATGATTTCGCGAAACTTGAGGAGCAGTTTGACGCTAAATTAGGCA<br>TTTTCGCCCTTGACACGGGAACCAACTCGACTGTCGCGTATCGCCCCGACGAGAGATTTGCCTTT<br>GCATCGACGATAAAAGCCTTGACAGTGGGGGTTCTGCTTCAACAAAAATCAATAGAGGACCTGAA<br>CCAACGTATTACCTACACAAGAGATGATTTAGTTAATTATAACCCAATAACAGAAAAACATGTGG<br>ACACAGGAATGACTTTGAAGGAATTGGCCGATGCAAGTTTAAGATACTCTGATAACGCCGCCCAA<br>AACCTGATCTTAAAACAGATCGGAGGCCCAGAGTCTTTGAAGAAAGAATTACGCAAAATAGGTGA<br>CGAGGTTACGAATCCCGAACGCTTCGAGCCCGAATTGAACGAAGTCAATCCAGGTGAGACACAGG<br>ACACATCTACAGCCCGCGCTCTGGTTACCAGTCTTAGAGCCTTTGCCCTGGAGGATAAATTACCG<br>TCCGAAAAGAGAGAACTTCTGATAGATTGGATGAAGAGAAACACTACGGGAGACGCCCTTATCAG<br>AGCGGGCGTCCCCGATGGGTGGAGGTGGCAGACAAGACTGGTGCGGCCTCTTATGGTACAAGAA<br>ACGACATTGCCATCATATGGCCCCCTAAGGGAGACCCCGTCGTGTTAGCAGTGTTAAGCAGTCGG<br>GACAAAAAAGATGCCAAATATGACGATAAGCTGATAGCTGAGGCGACAAAGGTTGTGATGAAGTT<br>ATTGAGAGCGCGGTCTACGCTGCAAAAGGAAGTGCATGCAGCTAAATCTTTAGCAATCATAGTCG<br>GGCTTTTTGCGTTGTGCTGGCTTCCGCTTCACATCATTAATTGTTTCACTTTCTTTTGTCCTGAC<br>TGCTCACACGCCCCCTTATGGCTTATGTACCTGGCGATAGTGCTTTCTCATACCAACTCTGTGGT<br>AAATCCTTTCATCTACGCGTACCGTATTCGGGAATTCCGTCAAACTTTTAGAAAAATAATCCGCA<br>GCCACGTCTTACGTCAGCAGGAGCCTTTCAAAGCGGCTGGGACATCAGCCAGAGTATTGGCAGCT<br>CATGGCTCGGATGGGGAGCAAGTTTCATTAAGATTAAATGGGCATCCGCCTGGAGTGTGGGCTAA<br>CGGCTCAGCACCCCATCCGGAGCGCCGTCCAAATGGTTATGCATTAGGGCTGGTATCCGGTGGCT<br>CCGCGCAGGAAAGTCAAGGCAATACCGGATTACCCGACGTTGAGCTGCTGTCACACGAACTGAAA<br>GGCGTTTGTCCAGAACCACCAGGCTTAGACGACCCTCTTGCTCAAGATGGCGCAGGCGTTTCGGA<br>GAACCTGTACTTCCAGAGC |
| BLIP-II | O87916 | 40-311, C-term 6XHis | CTCGAGGTTCTGTTCCAAGGACCTGCTACCAGCGTGGTAGCATGGGGGGGGAACAACGATTGGGG<br>TGAGGCTACAGTTCCGGCAGAGGCGCAATCAGGTGTGGACGCCATTGCTGGTGGTTACTTTCACG<br>GATTGGCGCTGAAGGGTGGCAAAGTTCTGGGCTGGGGAGCCAACCTTAACGGACAGCTGACCATG<br>CCCGCCGCCACACAGTCCGGTGTAGACGCCATTGCAGCCGGTAACTATCATAGCTTAGCACTTAA<br>AGACGGCGAAGTTATCGCATGGGGAGGGAATGAGGATGGGCAGACGACCGTTCCAGCCGAAGCGC<br>GTAGCGGCGTAGATGCTATAGCAGCTGGGGCATGGGCGTCCTATGCCTTGAAAGACGGGAAGGTC<br>ATAGCGTGGGGTGACGATTCTGATGGGCAAACAACTGTCCCGGCGGAAGCTCAGAGTGGAGTAAC<br>AGCGTTGGATGGCGGAGTATATACCGCGTTAGCGGTGAAAAACGGTGGCGTGATCGCTTGGGGGG<br>ATAATTACTTTGGGCAAACTACAGTCCCTGCCGAGGCGCAAAGCGGAGTTGACGATGTAGCGGGT<br>GGGATTTTTCATTCCCTGGCGTTGAAAGACGGGAAAGTGATCGCGTGGGGGGATAATAGATACAA<br>ACAGACTACAGTTCCGACTGAAGCTTTATCGGGCGTTTCTGCTATTGCTAGCGGAGAGTGGTATA<br>GTTTGGCACTTAAAAATGGGAAGGTGATCGCTTGGGGAAGTAGTCGGACTGCTCCGAGTAGCGTG<br>CAGAGCGGTGTGTCGTCTATAGAGGCGGGTCCGAATGCTGCGTATGCCCTTAAGGGAGGTTCGGG<br>CAGTGGGCACCATCACCATCATCACCACCATCATCATGAGAACCTGTACTTCCAGAGC |
| CCR10 | P46092 | N-term Met removed | CTCGAGGTTCTGTTCCAAGGACCTGGTACTGAAGCGACCGAACAAGTTAGCTGGGGCCACTATTC<br>TGGAGATGAAGAAGATGCCTATAGCGCAGAGCCTCTGCCGGAACTTTGTTATAAAGCCGATGTTC<br>AGGCTTTCAGCCGTGCTTTTCAGCCGAGCGTGTCCCTGACCGTAGCAGCCTTGGGCTTAGCCGGC<br>AACGGTCTGGTCCTGGCAACCCATCTGGCCGCACGCCGTGCAGCCCGCTCCCCTACCTCAGCCCA<br>CTTGCTGCAGCTGGCTCTTGCCGATCTGCTGCTGGCACTGACTCTTCCGTTTGCGGCTGCGGGTG<br>CGTTACAGGGCTGGAGTTTGGGCTCCGCCACCTGTCGCACCATTTCCGGTCTTTACAGCGCAAGC<br>TTCCACGCCGGCTTTCTGTTCCTGGCCTGTATTAGCGCCGACCGTTACGTGGCAATCGCCCGCGC<br>CCTGCCGGCCGGCCCGCGCCCTAGCACCCCGGGCCGTGCTCATTTAGTGTCAGTTATTGTCTGGC<br>TTCTGTCCTTACTGTTGGCTTTACCGGCGCTGTTATTTAGTCAAGACGGTCAGCGTGAAGGTCAA<br>CGTCGCTGCCGCCTGATTTTTCCTGAAGGCCTGACTCAGACTGTTAAAGGGGCATCGGCAGTCGC<br>GCAGGTCGCACTGGGTTTTGCCCTGCCGCTTGGAGTAATGGTTGCCTGCTACGCCCTGCTGGGCC<br>GCACACTGTTGGCGGCCCGTGGGCCTGAACGCCGCCGCGCGCTGCGTGTGGTGGTAGCGTTGGTA<br>GCGGCCTTTGTGGTCCTTCAGCTGCCCTACAGCCTGGCCCTTCTGTTGGACACCGCAGATCTGCT<br>GGCGGCCCGTGAGCGTAGCTGCCCGGCTTCCAAACGCAAAGATGTAGCCCTGCTGGTGACCAGCG<br>GTCTGGCCTTAGCTCGTTGCGGCCTGAATCCAGTACTGTACGCATTCCTGGGCCTGCGTTTTCGC<br>CAGGATCTTCGTCGCCTGTTACGTGGTGGCAGTTGCCCAAGCGGTCCGCAACCGCGTCGTGGCTG<br>CCCGCGTCGCCCGCGCCTGTCGTCTTGCTCAGCGCCTACCGAAACTCATTCGCTTAGCTGGGACA<br>ATGAGAACCTGTACTTCCAGAGC |
| CXCR1 | P25024 | N-term Met | CTCGAGGTTCTGTTCCAAGGACCTTCGAATATCACGGATCCTCAAATGTGGGATTTCGATGACCT<br>GAATTTTACTGGGATGCCGCCTGCGGACGAGGACTATTCCCCCTGCATGCTGGAAACGGAGACTT |

| | | | |
|---|---|---|---|
| | | removed | TAAACAAGTATGTTGTAATTATAGCTTACGCGTTGGTTTTCCTGCTGAGCCTGCTGGGTAATTCT<br>TTGGTTATGCTTGTTATTTTATATTCCCGCGTAGGCCGTTCCGTTACGGACGTTTATCTGCTGAA<br>TCTGGCCTTAGCTGACTTGCTGTTCGCTTTGACATTGCCAATCTGGGCCGCTAGCAAAGTAAACG<br>GTTGGATATTCGGAACCTTTTTATGCAAGGTGGTTTCCCTGCTTAAAGAAGTGAACTTTTACAGT<br>GGGATACTGCTGCTTGCTTGTATTTCTGTAGATCGCTACCTGGCGATTGTTCACGCTACTAGAAC<br>CCTGACGCAAAAACGTCATCTTGTAAAGTTCGTATGCCTGGGGTGCTGGGGATTATCGATGAATT<br>TATCGTTACCGTTCTTTCTGTTTAGACAGGCCTACCACCCCAATAATTCTTCCCCCGTCTGTTAT<br>GAGGTCCTGGGTAACGATACTGCAAAGTGGCGTATGGTCTTGCGCATTCTTCCCCATACTTTCGG<br>CTTCATTGTGCCGTTATTTGTGATGTTGTTTTGTTATGGCTTCACCTTACGGACGTTATTCAAGG<br>CGCATATGGGTCAGAAGCATCGTGCTATGAGAGTGATTTTCGCAGTCGTACTGATCTTCCTGCTT<br>TGTTGGTTACCATACAACCTGGTGCTTCTGGCTGACACGCTTATGAGAACGCAGGTTATCCAAGA<br>GAGCTGCGAAAGACGTAACAACATAGGGAGAGCGTTGGATGCGACTGAGATTTTAGGGTTCCTTC<br>ACTCATGTCTGAATCCAATAATTTATGCTTTTATAGGTCAGAATTTTCGGCACGGCTTCTTGAAA<br>ATTCTGGCTATGCACGGCCTGGTTTCCAAGGAATTTTTAGCGCGTCACCGGGTGACCAGTTACAC<br>CTCTAGCTCTGTTAATGTGTCCTCAAATTTGGAGAACCTGTACTTCCAGAGC |
| ADRB2 | P07550 | N-term Met removed | CTCGAGGTTCTGTTCCAAGGACCTGGGCAACCAGGTAATGGATCGGCATTCTTACTTGCGCCGAA<br>TGGAAGCCATGCACCGGATCACGATGTAACCCAGGAACGGGACGAGGTCTGGGTGGTGGGTATGG<br>GTATAGTAATGAGTCTGATAGTCCTTGCGATCGTCTTCGGCAACGTACTGGTTATCACTGCGATT<br>GCGAAGTTTGAACGCTTACAGACGGTGACCAACTACTTTATTACATCTCTTGCATGCGCGGATTT<br>AGTAATGGGATTAGCCGTCGTGCCGTTTGGAGCAGCACACATTCTTATGAAGATGTGGACGTTTG<br>GTAACTTTTGGTGTGAGTTCTGGACCTCCATTGACGTGTTGTGCGTGACGGCATCAATTGAAACC<br>CTGTGTGTGATCGCCGTCGATCGTTATTTTGCCATCACTAGTCCGTTTAAATATCAGAGTTTGTT<br>GACCAAAAACAAAGCTCGTGTAATCATTCTTATGGTTTGGATAGTCTCGGGTTTAACTTCGTTTT<br>TGCCAATACAGATGCATTGGTATCGCGCTACTCATCAGGAAGCCATCAATTGCTACGCCAACGAG<br>ACTTGCTGCGACTTTTTCACCAACCAGGCATATGCCATTGCCTCCTCGATCGTATCTTTTTACGT<br>CCCGTTGGTCATCATGGTGTTCGTCTACAGTCGGGTTTTTCAGGAGGCCAAAAGACAATTACAAA<br>AGATTGACAAAAGCGAGGGCCGTTTTCACGTGCAGAACTTATCCCAGGTCGAGCAAGATGGCCGG<br>ACAGGGCATGGCTTACGCCGCTCCTCAAAATTCTGCCTTAAGGAACATAAAGCCCTTAAGACTCT<br>TGGTATAATTATGGGGACATTTACCCTGTGTTGGCTGCCCTTCTTTATAGTGAACATAGTGCACG<br>TTATCCAGGACAATCTGATCCGGAAAGAAGTATACATACTTCTGAATTGGATCGGGTATGTGAAT<br>TCTGGATTCAACCCTCTGATTTATTGCAGATCTCCGGATTTTAGAATTGCTTTTCAAGAGCTTTT<br>GTGCCTGCGTCGTAGCAGTTTGAAGGCTTATGGTAACGGCTATTCTTCTAACGGCAATACAGGAG<br>AACAAAGTGGCTACCATGTGGAGCAAGAAAAGGAGAATAAATTATTGTGTGAAGACCTGCCGGGG<br>ACCGAGGACTTTGTGGGGCACCAAGGCACAGTTCCTTCAGATAATATAGATTCTCAGGGTCGTAA<br>CTGCTCCACCAATGACTCACTTTTAGAGAACCTGTACTTCCAGAGC |
| CB2 | P34972 | N-term Met removed | CTCGAGGTTCTGTTCCAAGGACCTGAAGAGTGTTGGGTTACAGAAATTGCCAACGGCTCGAAAGA<br>TGGCCTGGACAGCAATCCGATGAAAGACTACATGATTTTGAGCGGGCCGCAGAAAACGGCAGTGG<br>CCGTCCTTTGCACCCTGTTAGGTCTGCTGTCAGCCCTTGAAAATGTCGCGGTTTTGTATCTTATC<br>CTGAGCAGCCATCAGTTACGCCGCAAACCCTCATATCTGTTTATCGGAAGTCTGGCAGGGGCCGA<br>TTTTCTTGCGTCAGTGGTATTCGCCTGCTCATTCGTCAACTTCCATGTGTTTCATGGTGTTGATT<br>CGAAAGCCGTATTTCTTTTGAAAATTGGCTCTGTGACCATGACCTTCACCGCAAGCGTAGGCTCA<br>TTACTGTTAACTGCAATCGATCGTTATTTATGTCTGCGTTACCCTCCGAGCTATAAAGCGCTGCT<br>GACACGCGGTCGTGCGCTTGTTACCCTGGGAATCATGTGGGTACTTTCGGCGCTTGTGAGCTATC<br>TTCCGCTGATGGGTTGGACGTGTTGCCCGCGCCCGTGCTCTGAACTGTTTCCGCTGATCCCGAAT<br>GACTACTTACTGAGCTGGTTATTATTCATTGCGTTTCTGTTTAGCGGTATCATTTACACGTACGG<br>TCACGTACTGTGGAAAGCGCATCAGCACGTGGCCTCGCTGTCGGGTCATCAAGACCGCCAGGTTC<br>CAGGTATGGCACGCATGCGCTTAGATGTGCGCTTGGCGAAAACTCTGGGGCTGGTGTTGGCGGTC<br>CTGCTGATCTGTTGGTTCCCCGTGCTGGCTCTTATGGCCCATTCACTTGCCACGACCCTGAGCGA<br>TCAGGTCAAAAAGGCGTTTGCCTTTTGCAGCATGTTGTGCTTGATCAACTCTATGGTGAACCCCG<br>TCATTTACGCGTTGCGCTCTGGTGAAATTCGTTCCAGCGCACATCATTGCCTGGCACACTGGAAA<br>AAGTGTGTACGCGGTCTGGGTTCTGAAGCTAAAGAGGAAGCTCCACGCTCGTCTGTAACGGAAAC<br>GGAAGCAGATGGAAAAATCACCCCGTGGCCGGATAGCCGCGACCTGGATTTGAGCGATTGTGAGA<br>ACCTGTACTTCCAGAGC |
| GCGR | P47871 | N-term Met removed | CTCGAGGTTCTGTTCCAAGGACCTGCCCAGGTAATGGATTTCCTGTTTGAAAAATGGAAGCTGTA<br>CGGCGATCAGTGTCATCATAATTTATCCCTGCTGCCACCACCGACCGAGCTGGTCTGTAATCGTA<br>CCTTTGATAAATATAGCTGTTGGCCGGACACACCTGCGAACACGACCGCAAACATTTCGTGTCCC<br>TGGTATCTGCCGTGGCACCATAAGGTGCAGCATCGTTTTGTGTTCAAACGCTGCGGCCCTGATGG<br>GCAGTGGGTCCGTGGTCCGCGCGGTCAGCCGTGGCGCGATGCAAGCCAGTGCCAAATGGATGGTG<br>AGGAAATTGAAGTGCAGAAGGAGGTCGCGAAGATGTATAGCAGCTTCCAAGTGATGTACACGGTT<br>GGGTACTCCCTGAGCCTGGGCGCGCTGCTGCTGGCGCTGGCGATTCTGGGTGGACTGAGTAAGCT |

| | | | |
|---|---|---|---|
| | | | GCATTGTACTCGTAATGCCATTCACGCAAATCTGTTTGCATCGTTCGTTCTTAAGGCCAGCAGTG<br>TGCTGGTCATTGATGGTTTACTGCGTACCCGTTATAGCCAGAAAATTGGTGATGATCTGTCTGTT<br>TCCACTTGGTTATCTGATGGGGCCGTTGCAGGATGTCGCGTAGCGGCAGTTTTCATGCAGTATGG<br>CATTGTGGCCAACTATTGTTGGCTTTTGGTTGAAGGTTTGTACTTGCACAACTTGCTGGGTCTTG<br>CCACCCTGCCTGAACGTAGTTTTTTCTCACTGTACTTAGGCATCGGTTGGGGCGCGCCAATGCTG<br>TTTGTCGTGCCGTGGGCGGTCGTCAAATGCCTGTTTGAAAACGTGCAGTGCTGGACCAGCAATGA<br>CAACATGGGCTTCTGGTGGATCTTACGTTTCCCTGTTTTCTTAGCCATCTTAATCAATTTTTTTA<br>TTTTTGTACGTATTGTGCAGCTTTTAGTAGCCAAGTTACGCGCGCGTCAGATGCACCACACCGAT<br>TACAAATTTCGTCTGGCAAAGTCGACGCTTACGCTTATTCCGCTGCTGGGCGTGCACGAGGTTGT<br>CTTTGCGTTTGTGACGGATGAACATGCGCAGGGGACCCTGCGTAGCGCAAAACTGTTTTTCGATT<br>TGTTTCTTTCGAGCTTCCAGGGGTTACTGGTAGCCGTTCTGTATTGTTTTCTGAATAAAGAGGTG<br>CAGTCGGAACTGCGCCGCCGCTGGCATCGTTGGCGCTTAGGTAAAGTGCTGTGGGAAGAACGCAA<br>CACGAGCAATCATCGTGCCTCCAGCTCGCCTGGCCACGGTCCCCCTTCAAAAGAATTGCAGTTTG<br>GTCGCGGGGGTGGCTCACAAGATAGCAGTGCGGAAACCCCCCTGGCAGGCGGCCTGCCTCGCCTT<br>GCGGAATCGCCATTTGAGAACCTGTACTTCCAGAGC |
| RAI3 | Q8NFJ5 | N-term Met removed | CTCGAGGTTCTGTTCCAAGGACCTGCCACCACCGTGCCCGACGGTTGTCGCAATGGTTTGAAGTC<br>AAAATACTATCGGCTGTGTGATAAAGCGGAGGCTTGGGGAATCGTACTGGAAACGGTTGCGACGG<br>CTGGAGTGGTGACATCAGTAGCATTTATGTTGACCTTGCCCATCCTTGTGTGCAAGGTTCAGGAC<br>AGTAATCGCAGAAAGATGTTACCAACCCAATTTCTTTTCTTGCTGGGCGTATTGGGTATATTTGG<br>ACTGACATTCGCGTTTATCATTGGGTTAGATGGCTCAACCGGGCCCACTAGATTTTTCCTTTTCG<br>GTATACTGTTTAGTATCTGTTTTTCGTGTTTACTGGCTCATGCAGTATCGTTAACTAAATTAGTC<br>CGGGGTCGTAAGCCGCTTTCTCTGTTGGTTATACTGGGTTTAGCGGTTGGATTTAGCTTGGTGCA<br>GGATGTGATTGCGATAGAGTACATAGTTCTGACAATGAATCGTACAAACGTGAATGTATTCTCAG<br>AATTATCGGCACCACGTCGGAACGAGGATTTTGTTTTACTGCTTACGTATGTATTATTTTTGATG<br>GCGCTTACTTTTCTTATGAGCTCCTTCACGTTCTGCGGGTCGTTTACAGGTTGGAAACGTCACGG<br>GGCTCATATTTATTTGACTATGCTTCTTAGTATCGCAATATGGGTCGCGTGGATAACTTTGCTTA<br>TGTTACCTGATTTCGACAGAAGATGGGACGACACTATACTTAGCTCGGCCTTGGCCGCTAACGGC<br>TGGGTCTTCCTGTTGGCTTATGTTTCTCCTGAGTTTTGGTTGCTTACCAAACAAAGAAACCCCAT<br>GGACTATCCGGTCGAAGACGCATTCTGTAAGCCACAGTTAGTAAAAAAATCGTATGGCGTCGAGA<br>ATAGAGCGTATAGCCAGGAAGAGATCACCCAGGGCTTTGAGGAAACCGGTGATACTTTGTATGCA<br>CCCTATAGCACTCACTTCCAACTGCAAAATCAGCCGCCTCAAAAAGAATTTTCAATACCGAGAGC<br>TCACGCTTGGCCGTCGCCTTACAAGGACTACGAAGTGAAGAAAGAAGGATCAGAGAACCTGTACT<br>TCCAGAGC |
| GPR61 | Q9BZJ8 | N-term Met removed | CTCGAGGTTCTGTTCCAAGGACCTGAATCATCTCCGATCCCTCAGTCGTCTGGTAATAGCAGCAC<br>GCTGGGACGCGTCCCGCAGACCCCAGGCCCTAGTACCGCCAGCGGCGTACCAGAAGTCGGCCTGC<br>GCGACGTTGCAAGTGAAAGCGTTGCCCTGTTTTTCATGCTGTTGTTAGACCTGACGGCAGTCGCC<br>GGGAATGCCGCGGTTATGGCGGTCATTGCGAAAACCCCAGCGCTGCGCAAGTTCGTCTTTGTGTT<br>TCACCTTTGCCTGGTGGACTTGCTTGCCGCGCTGACACTGATGCCGCTGGCCATGCTGAGCTCGA<br>GCGCCCTGTTTGATCACGCTCTGTTTGGTGAGGTTGCCTGCCGCCTTTACCTTTTTCTGAGCGTT<br>TGCTTTGTTTCGTTAGCCATTCTTTCGGTGTCTGCGATTAACGTAGAACGCTATTACTACGTGGT<br>ACACCCTATGCGCTACGAAGTTCGTATGACGCTGGGTCTGGTTGCGTCTGTGTTAGTCGGCGTGT<br>GGGTCAAAGCCTTAGCGATGGCGTCCGTTCCGGTTTTAGGCCGCGTTTCGTGGGAAGAGGGCGCC<br>CCGAGCGTCCCTCCGGGTTGCAGCCTGCAGTGGTCCCATTCCGCCTACTGCCAACTGTTTGTGGT<br>AGTGTTTGCAGTGCTGTACTTCCTGCTGCCTCTGCTGTTGATCCTGGTTGTGTACTGCAGCATGT<br>TCCGCGTGGCTCGTGTTGCCGCCATGCAGCATGGACCGTTGCCGACGTGGATGGAGACCCCGCGT<br>CAGCGCTCGGAGAGCCTGTCAAGTCGTAGCACCATGGTGACCAGCTCGGGTGCACCTCAAACCAC<br>CCCCCATCGTACCTTTGGTGGCGGAAAAGCCGCAGTTGTCCTGTTAGCCGTCGGTGGGCAGTTCC<br>TGTTGTGCTGGCTGCCGTATTTCTCTTTCCACCTGTACGTGGCGCTGAGCGCCCAGCCAATTTCT<br>ACTGGCCAGGTTGAAAGCGTCGTGACCTGGATTGGCTACTTTTGTTTTACAAGTAATCCCTTCTT<br>CTACGGATGCCTGAACCGTCAGATTCGTGGGGAACTGTCCAAACAGTTCGTATGCTTCTTCAAAC<br>CGGCGCCGGAAGAAGAATTACGCCTTCCGAGCCGCGAAGGCTCGATTGAAGAAAACTTCCTGCAA<br>TTTCTGCAGGGAACCGGATGTCCGAGCGAAAGTTGGGTAAGTCGCCCGCTTCCCTCGCCGAAACA<br>GGAGCCTCCGGCCGTAGATTTCCGCATTCCGGGACAGATCGCAGAAGAAACGAGCGAATTTCTTG<br>AACAGCAGCTGACCTCAGATATTATTATGTCGGATTCCTACTTACGTCCGGCCGCAAGCCCCCGC<br>TTGGAAAGCGAGAACCTGTACTTCCAGAGC |
| GPR85 | P60893 | N-term Met removed | CTCGAGGTTCTGTTCCAAGGACCTGCCAATTACAGTCACGCCGCCGATAATATTCTGCAGAACCT<br>GAGTCCTTTAACGGCGTTCCTGAAACTGACCTCACTGGGCTTCATCATTGGTGTGTCAGTTGTCG<br>GCAACTTGTTAATTAGTATCCTGCTGGTTAAAGATAAAACACTGCATCGCGCACCTTACTATTTT<br>CTGCTGGACCTGTGCTGCTCTGACATTTTACGCTCTGCAATTTGCTTCCCTTTCGTATTTAATTC<br>CGTTAAAAACGGTAGCACCTGGACATACGGTACCTTGACGTGCAAAGTTATTGCGTTTCTGGGTG |

| | | | |
|---|---|---|---|
| | | | TGCTTAGCTGCTTCCATACCGCGTTCATGCTGTTTTGTATTTCCGTCACCCGTTACTTAGCAATT<br>GCCCATCACCGTTTCTATACCAAACGTCTTACCTTTTGGACATGTCTTGCAGTAATTTGCATGGT<br>CTGGACGCTGAGCGTTGCCATGGCGTTTCCGCCGGTACTGGATGTGGGCACCTACAGCTTTATTC<br>GTGAGGAAGACCAATGCACGTTCCAGCATCGTAGCTTCCGCGCGAATGATTCTCTGGGCTTTATG<br>TTATTATTGGCGTTGATCTTGCTGGCGACTCAGCTGGTCTATCTGAAGCTGATTTTTTTCGTGCA<br>TGATCGCCGTAAGATGAAACCGGTGCAGTTCGTGGCGGCAGTGAGCCAGAACTGGACCTTCCACG<br>GGCCTGGAGCCAGCGGCCAGGCCGCAGCGAATTGGCTGGCCGGTTTTGGACGTGGCCCCACTCCA<br>CCTACCTTACTGGGCATCCGCCAGAATGCCAACACGACCGGTCGCCGCCGTCTGCTTGTGCTTGA<br>TGAATTTAAGATGGAGAAGCGCATCAGCCGTATGTTCTACATCATGACCTTTCTGTTTCTTACCC<br>TGTGGGGTCCGTATCTGGTAGCGTGTTACTGGCGCGTGTTTGCGCGTGGCCCTGTCGTCCCGGGC<br>GGCTTCTTAACAGCCGCAGTTTGGATGTCATTTGCACAGGCCGGCATCAATCCCTTTGTGTGTAT<br>TTTCAGCAACCGCGAGCTGCGCCGTTGCTTTAGCACCACGTTATTATATTGCCGCAAGTCGCGTC<br>TTCCGCGTGAACCCTACTGTGTCATCGAGAACCTGTACTTCCAGAGC |
| GPR88 | Q9GZN0 | N-term Met removed | CTCGAGGTTCTGTTCCAAGGACCTACTAACTCGTCATCCACCTCCACGAGCTCCACCACCGGCGG<br>CTCTCTGCTTCTGCTTTGTGAAGAAGAATCGTGGGCTGGACGTCGCATCCCCGTTAGCTTGC<br>TTTATAGCGGTCTGGCGATTGGTGGCACGCTGGCAAACGGCATGGTGATTTACCTGGTGTCGTCT<br>TTTCGCAAACTGCAGACAACGTCGAACGCATTTATTGTGAATGGTTGCGCGGCTGATCTTTCGGT<br>CTGCGCGTTATGGATGCCTCAAGAAGCAGTACTGGGGCTGCTGCCGACCGGCAGTGCGGAACCGC<br>CGGCGGACTGGGACGGTGCCGGTGGTTCGTACCGCCTGTTGCGCGGCGGGTTATTGGGGTTAGGG<br>CTGACTGTCTCTCTGCTGTCACACTGCTTAGTGGCGTTAAACCGCTACCTGCTGATTACGCGCGC<br>CCCGGCCACCTACCAGGCGTTGTATCAGCGCCGCCACACCGCAGGCATGCTGGCATTATCGTGGG<br>CGCTTGCATTAGGATTAGTGCTGTTGCTTCCGCCATGGGCGCCTCGCCCTGGTGCCGCCCCGCCG<br>CGCGTGCATTATCCGGCACTGCTTGCCGCGGCCGCGCTGCTGGCCCAAACAGCACTTCTGCTGCA<br>CTGCTATTTAGGTATTGTCCGCCGCGTGCGCGTTTCAGTTAAACGCGTGTCCGTGCTTAACTTCC<br>ACCTTCTGCACCAATTACCAGGCTGCGCGGCAGCAGCAGCCGCGTTCCCGGGTGCGCAGCATGCG<br>CCGGGACCGGGCGGAGCGGCACATCCGGCACAAGCCCAGCCATTGCCGCCCGCCCTGCACCCGCG<br>CCGTGCACAACGTCGCCTGTCGGGTTTAAGCGTCTTGCTGCTGTGTTGCGTCTTTTTACTTGCGA<br>CACAACCGCTGGTTTGGGTGTCTCTTGCGTCTGGCTTTAGCTTACCGGTGCCGTGGGGCGTGCAA<br>GCCGCCAGCTGGCTGCTGTGTTGCGCCCTGAGCGCACTTAACCCGTTACTGTATACTTGGCGCAA<br>CGAGGAATTTCGTCGTAGCGTGCGTAGTGTGCTTCCTGGCGTCGGGGATGCGGCAGCAGCCGCAG<br>TGGCCGCGACTGCCGTACCAGCCGTTTCGCAGGCACAACTGGGTACCCGCGCAGCGGGTCAGCAT<br>TGGGAGAACCTGTACTTCCAGAGC |
| GPR146 | Q96CH1 | N-term Met removed | CTCGAGGTTCTGTTCCAAGGACCTTGGAGTTGTTCGTGGTTCAACGGCACGGGATTAGTCGAAGA<br>ACTTCCTGCCTGCCAGGATCTGCAACTTGGATTATCTCTGCTGTCACTGTTAGGCCTGGTTGTTG<br>GCGTGCCGGTAGGTCTGTGCTACAATGCGCTGCTGGTGTTGGCCAACCTGCATTCCAAGGCGAGT<br>ATGACAATGCCGGATGTCTACTTTGTAAATATGGCCGTGGCGGGCTTGGTGCTGAGCGCTTTAGC<br>CCCAGTGCACTTGCTGGGCCCTCCGTCAAGCCGTTGGGCTTTATGGTCGGTAGGGGGAGAAGTTC<br>ACGTCGCCCTGCAGATTCCGTTCAATGTGAGCAGCCTGGTAGCGATGTATTCAACCGCGTTGCTG<br>TCGCTGGATCATTATATCGAACGCGCACTGCCGCGCACCTATATGGCGAGTGTGTACAACACTCG<br>TCATGTATGTGGCTTTGTATGGGGCGGGGCCCTGCTGACATCCTTTAGCTCTCTGCTGTTTTACA<br>TCTGCAGCCACGTCAGCACTCGCGCCTTAGAGTGTGCGAAAATGCAGAACGCAGAAGCGGCTGAT<br>GCTACCCTGGTGTTTATTGGCTACGTCGTGCCTGCGCTGGCGACCCTGTACGCCCTGGTGCTGTT<br>GAGCCGTGTACGCCGCGAGGATACCCCGTTGGATCGCGATACCGGGCGCTTGGAACCATCCGCAC<br>ATCGCCTGTTGGTGGCGACTGTCTGCACGCAGTTTGGGCTGTGGACCCCGCACTATCTGATTTTA<br>CTGGGCCACACTGTGATTATTAGCCGTGGTAAACCTGTTGACGCCCATTACTTGGGCCTGCTGCA<br>CTTCGTTAAAGATTTCTCAAAACTTTTGGCTTTTTCATCCTCATTTGTTACTCCTCTTTTATACC<br>GTTACATGAACCAGTCATTCCCGTCAAAACTGCAGCGCCTGATGAAAAAACTGCCCTGTGGTGAC<br>CGCCACTGTTCTCCCGACCATATGGGTGTGCAGCAGGTGCTGGCGGAGAACCTGTACTTCCAGAG<br>C |
| GPR151 | Q8TDV0 | N-term Met removed | CTCGAGGTTCTGTTCCAAGGACCTTTGGCAGCGGCATTTGCCGATAGCAATAGCTCGTCAATGAA<br>CGTGAGCTTTGCACATTTACATTTCGCTGGGGGGTACCTGCCGAGTGATAGCCAGGACTGGCGCA<br>CAATTATCCCGGCATTACTTGTTGCTGTCTGTTTAGTAGGCTTTGTCGGTAATCTTTGTGTAATT<br>GGGATCCTGCTGCATAATGCATGGAAAGGCAAACCGTCGATGATCCATAGTCTTATCCTGAACCT<br>GTCTTTAGCGGACCTGTCGCTGCTGCTGTTTAGCGCACCGATTCGTGCAACCGCGTACAGCAAAT<br>CGGTGTGGGACCTGGGGTGGTTTGTATGCAAAAGCAGTGACTGGTTCATCCACACGTGCATGGCG<br>GCGAAGTCATTAACGATCGTAGTCGTCGCAAAGGTGTGCTTTATGTATGCATCGGATCCGGCGAA<br>GCAAGTCTCAATTCACAATTATACGATTTGGAGTGTGCTTGTAGCGATCTGGACCGTGGCGTCGT<br>TGCTGCCTCTGCCGGAATGGTTCTTTTCAACGATTCGCCATCACGAGGGCGTGGAAATGTGCCTT<br>GTGGATGTGCCTGCCGTAGCTGAAGAGTTCATGTCAATGTTTGGCAAACTGTACCCATTGCTGGC<br>GTTCGGCCTGCCACTGTTTTTTGCGAGCTTCTATTTTTGGCGTGCATACGACCAGTGTAAAAAAC |

| | | | |
|---|---|---|---|
| | | | GCGGAACTAAAACCCAGAATCTTCGCAACCAGATCCGTAGCAAACAGGTCACGGTTATGCTGCTG<br>AGTATCGCGATCATTTCAGCCTTATTATGGTTGCCTGAATGGGTGGCATGGCTGTGGGTATGGCA<br>TCTGAAGGCAGCGGGTCCGGCTCCTCCGCAGGGCTTCATCGCCCTTAGTCAAGTCCTGATGTTTA<br>GCATTAGCAGCGCGAACCCGCTGATCTTTTTGGTTATGTCCGAAGAATTCCGCGAAGGCTTAAAA<br>GGAGTGTGGAAATGGATGATCACCAAGAAGCCTCCGACCGTATCCGAATCTCAGGAAACACCGGC<br>TGGCAATAGTGAAGGGCTGCCGGACAAGGTCCCTAGCCCGGAGAGCCCAGCATCCATTCCGGAAA<br>AAGAAAAGCCATCCAGCCCGTCATCGGGGAAAGGCAAGACTGAAAAGGCCGAAATTCCAATCTTG<br>CCGGATGTAGAACAGTTCTGGCACGAGCGCGATACCGTCCCTTCCGTGCAAGATAATGACCCGAT<br>TCCGTGGGAGCACGAAGACCAAGAAACTGGAGAAGGCGTTAAAGAGAACCTGTACTTCCAGAGC |
| GPR174 | Q9BXC1 | N-term Met removed | CTCGAGGTTCTGTTCCAAGGACCTCCGGCGAACTACACCTGCACGCGCCCGGATGGAGATAACAC<br>TGATTTCCGCTATTTCATTTACGCGGTTACTTACACGGTGATTTTAGTGCCGGGGTTAATTGGTA<br>ACATTTTAGCGCTGTGGGTTTTCTATGGTTATATGAAAGAGACCAAACGTGCGGTTATTTTTATG<br>ATTAATCTGGCCATTGCGGATTTATTACAGGTCCTGAGTTTACCACTGCGTATCTTCTACTATCT<br>GAATCATGACTGGCCCTTTGGTCCCGGTCTGTGCATGTTCTGCTTCTACCTGAAATACGTGAACA<br>TGTATGCGAGTATCTATTTTCTGGTGTGTATTAGCGTCCGTCGTTTCTGGTTTCTGATGTATCCG<br>TTTCGTTTTCATGATTGCAAACAGAAATATGATCTGTACATCAGTATTGCGGGCTGGCTTATTAT<br>TTGTCTGGCTTGCGTGCTGTTCCCATTACTGCGTACCTCAGATGATACAAGCGGTAATCGCACCA<br>AGTGCTTCGTTGACCTGCCAACCCGCAACGTCAATCTGGCGCAAAGCGTTGTTATGATGACCATT<br>GGCGAGCTGATTGGTTTTGTGACCCCGCTGCTGATTGTTTTATATTGCACCTGGAAGACCGTGCT<br>GTCCCTTCAGGATAAGTACCCGATGGCGCAAGACTTGGGGGAGAAGCAAAAAGCGCTGAAGATGA<br>TCCTGACTTGCGCCGGCGTTTTTCTGATTTGTTTTGCGCCGTATCATTTTTCGTTCCCTTTAGAT<br>TTTCTTGTTAAAAGCAATGAAATCAAATCATGTCTGGCCCGTCGTGTGATTCTGATTTTCCACTC<br>TGTTGCCCTGTGCCTGGCCTCACTGAATAGCTGTCTGGACCCGGTTATCTATTACTTTTCGACCA<br>ATGAGTTCCGCCGCCGTCTGTCTCGTCAGGACCTTCACGATTCTATTCAACTTCACGCTAAGTCG<br>TTCGTGTCAAATCACACGGCGAGCACGATGACGCCGGAGTTATGTGAGAACCTGTACTTCCAGAG<br>C |

***Table S2.*** *gBlock sequences ordered for eGene assembly and cell-free expression. Underlined sequences indicate 3C and TEV cleavage sites used for eGene assembly.*

| | PDB: xxxx<br>EMD: xxxx |
|---|---|
| **Data collection and processing** | |
| Detector | Falcon 4i |
| Magnification | 165k |
| Energy filter slit width (eV) | 10 |
| Voltage (kV) | 300 |
| Flux on detector (e/pix/sec) | 8.36 |
| Electron exposure on sample (e–/Å$^2$) | 46.09 |
| Target defocus range (µm) | 0.6-1.8 |
| Calibrated pixel size (Å) | 0.729 |
| Symmetry imposed | C1 |
| Extraction box size (pixels) | 384 |
| Initial particle images (no.) | 5479759 |
| Final particle images (no.) | 47986 |
| **Refinement** | |
| Map resolution at FSC=0.143 (Å)* | 3.38 |
| Model composition | |
| Non-hydrogen atoms | 6115 |
| Protein residues | 803 |
| B factor (Å$^2$) | |
| Protein | 168.83 |
| R.m.s deviations | |
| Bond lengths (Å) | 0.004 |
| Bond angles (°) | 0.650 |
| Validation | |
| Molprobity score | 2.63 |
| Clashscore | 12.83 |
| Poor rotamers (%) | 3.79 |
| Ramachandran plot | |
| Favored (%) | 89.21 |
| Allowed (%) | 9.91 |
| Disallowed (%) | 0.88 |

***Supplementary Table 3.*** *Cryo EM data collection parameters.*

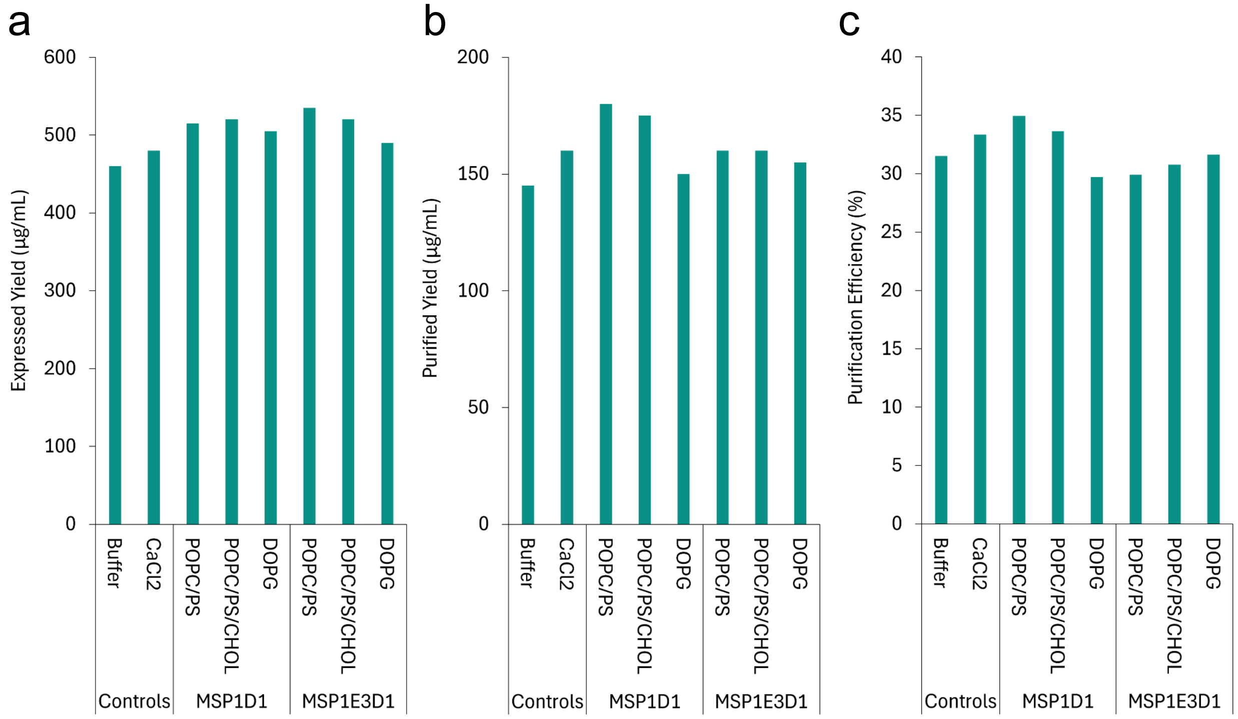


***Supplementary Figure 1.*** *Cell-free expression yields (a), purification yields (b), and purification efficiency (c) of BLIP-II.*

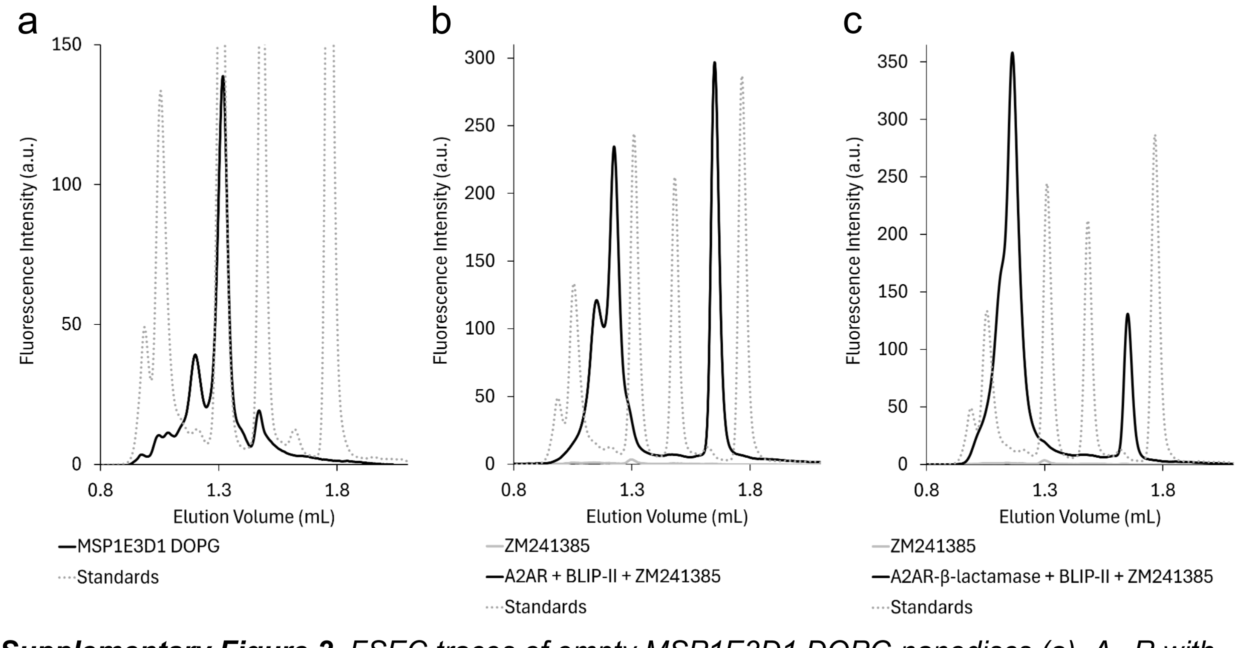


***Supplementary Figure 2.*** *FSEC traces of empty MSP1E3D1 DOPG nanodiscs (a), $A_{2A}R$ with BLIP-II and ZM241385 (b), and $A_{2A}R$-β-lactamase with BLIP-II and ZM241385 (c). Dotted gray traces indicate molecular weight standards (Waters, BEH200 SEC Test Mix) corresponding to 660 kDa, 150 kDa, 66.4 kDa, and 17 kDa.*

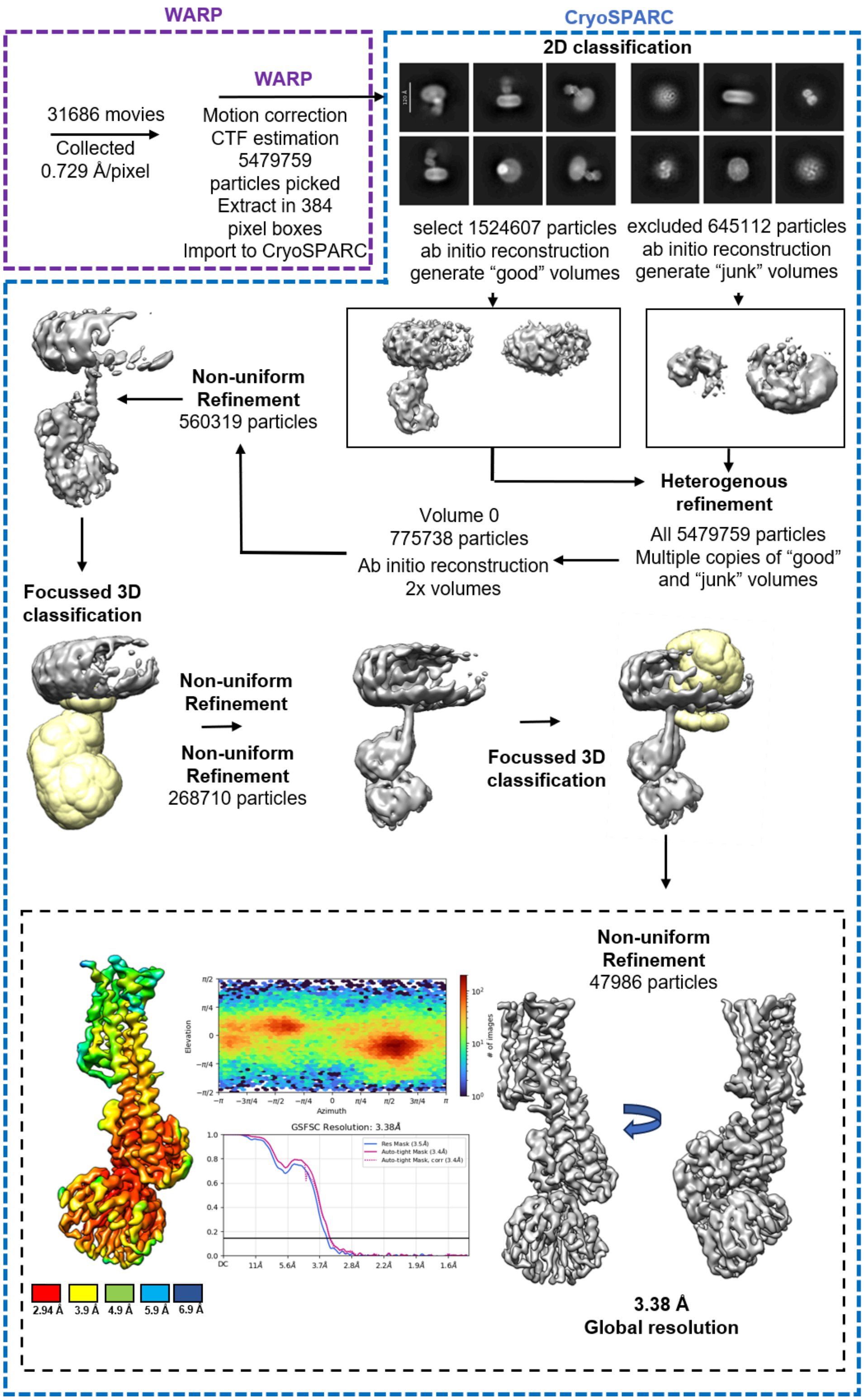


***Supplementary Figure 3.*** *Cryo-EM data processing workflow for $A_{2A}$R-β-lactamase in Complex with BLIP-II in MSP1E3D1 DOPG Nanodiscs. Upper panels show individual processing steps. Bottom panel (dashed black box) summarises final map quality. Left – local resolution map. Middle top, particle orientation plot. Middle bottom, FSC curves.*

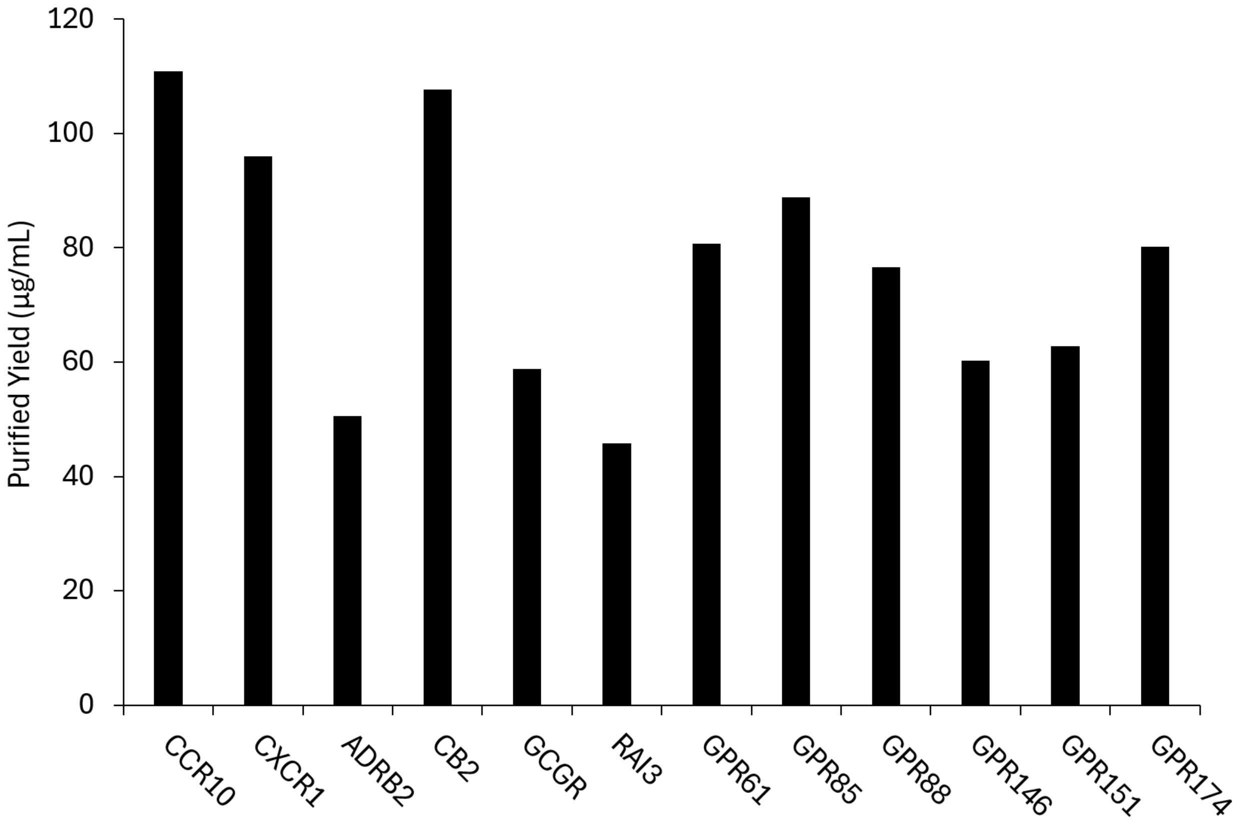


***Supplementary Figure 4.*** *Purification yields following scaled-up expression and purification of the expanded 12 GPCR panel. Expression conditions are shown in Table S1. FSEC analysis is shown in Figure 4.*
